# GmEBS7a-E2 complex dissociation hyperactivates ERAD to impair plant immunity against *Phytophthora* pathogens

**DOI:** 10.64898/2026.01.25.701614

**Authors:** Guangda Shao, Qingyu Liu, Xi Zhang, Hua Zhao, Wenxin Gao, Song Tian, Zeqi Liu, Qinhu Wang, Biao Gu, Xili Liu

## Abstract

Endoplasmic reticulum-associated degradation (ERAD) pathways are crucial for maintaining plant development and stress responses. While ERAD has been extensively studied for its role in regulating responses to abiotic stress, its specific manipulation by eukaryotic pathogens stress remains unclear. Here, we reported that soybean GmEBS7a, a plant-specific ERAD component, is targeted and stabilized by a transmembrane domain-containing effector Avh131 from *Phytophthora sojae*. GmEBS7a enhances plant disease susceptibility by increasing ERAD efficiency and ER stress sensitivity. Mechanistically, Avh131 competitively disrupts the interaction with GmEBS7a and the ubiquitin-conjugating enzyme GmUBC32, reducing ubiquitination of GmEBS7a and subsequent inhibition of its degradation. These findings demonstrate that excessive ERAD triggered by stabilized GmEBS7a and its partnering E3 ligase impairs the plant’s response to biotic stress. Our study uncovers a fine-tuning of the ER protein-folding capacity to promote infection, thereby expanding our understanding of ER-centered immunity during plant-pathogen interactions.

## Introduction

Plants have evolved sophisticated surveillance and defense systems to overcome diverse environmental stresses. Such sensing and responding systems depend mainly on coordinating different organelles and conserved cellular processes in eukaryotic cells. For example, the endoplasmic reticulum (ER) quality control machinery monitors the production and assembly of secreted and transmembrane domain (TMD)-containing proteins, many of which are well-known sensors and regulators against pathogen infection (*1*). Increasing evidence suggests that a plant’s ER and the regulation of its capacity play a crucial role in immunity (*2, 3*). Several plant pathogens can manipulate the ER stress response to benefit colonization and proliferation (*4*). Thus, the insight into stress-associated regulators of ER protein homeostasis will provide a better understanding of plant immune programming.

ER stress in plant cells is caused by the rapid accumulation or prolonged retention of unassembled, misfolded, or aggregated proteins in the ER. The consistent activation of ER stress-related mechanisms eventually leads to cell death; this can be an antagonistic strategy of the host plant to constrain pathogen infection (*4*), but it can also cause growth retardation (*5*). To mitigate the effects of ER stress, plants have evolved several distinct cytoprotective strategies, one of which is maintaining a dynamic protein equilibrium in the ER, known as ER-associated degradation (ERAD) (*5, 6*). Two conserved ER transmembrane ubiquitin E3 ligases, HMG-CoA reductase degradation 1 (Hrd1) and degradation of alpha2 10 (Doa10), can form a core complex involved in ERAD pathways in plants and are thus central modules for plant adaptation to both biotic and abiotic stresses (*7, 8*). An Hrd1a or Doa10 complex is required to degrade membrane-localized resistance protein and pattern recognition receptors (PRRs), such as MILDEW RESISTANCE LOCUS O (MLO) or ELONGATION FACTOR THERMO UNSTABLE RECEPTOR (EFR) (*9–11*). Plant-specific proteins can also be encoded to recognize different ERAD substrates.

AtEBS7, one such plant-specific ERAD component, interacts with and stabilizes AtHrd1a to reduce polyubiquitination of bri1-9 and regulate ER- and salt-based stresses (*11*). AtPAWH1 and AtPAWH2 are recently identified plant-specific components of ERAD; they maintain the stability of AtEBS7 and AtHrd1a and are essential for stress tolerance (*12*). A plant-specific ring finger E3 ligase (AtEMR) is associated with ubiquitin-conjugating enzyme 32 (UBC32) in the cytosol and is involved in the degradation of AtOS9 and mildew resistance locus O-12 (MLO12) (*13*). An ER-membrane-localized RING E3 (MfSTMIR) responds to ER stress and degrades its substrate and the vacuolar carboxypeptidase mutant MtCPY* (*14*). Although the functions of these ERAD components in plant development and the response to abiotic stress have been extensively documented, how ERAD machinery regulates plant immunity against fungal and oomycete pathogens remains to be explored.

Some ERAD components are involved in the activation and signal transduction of different plant immune responses. First, the ERAD pathway is tightly linked to pattern-triggered immunity (PTI). A membrane-bound leucine-rich-repeat receptor-like kinase, EFR, has been identified as an ERAD substrate (*15*). The Leucine-rich repeat kinase receptor (LRR-RLKs) co-receptor kinase SERK1 binds and phosphorylates the ATPase CDC48 (*16*), a key component of both Hrd1a and Doa10 complexes. MLO proteins are known to regulate basal resistance to certain biotrophic pathogens, with their ERAD-mediated degradation contributing to the modulation of PTI-related defenses (*9*). Second, the components of ERAD also regulate effector-triggered immunity (ETI). Coupling of ubiquitin conjugation to ER (CUE)-containing E3 ligases and *Arabidopsis* R protein RPM1, regulates the RPM1-and RPS2-dependent hypersensitive response (HR) (*17*). Silencing of ER luminal binding protein-coding genes *BiP4* and *BiP5* attenuates resistance protein HRT-mediated cell death (*18*). Additionally, plants often defend themselves against pathogens by inducing ER stress. For example, *Arabidopsis* RTP1 negatively regulates the activation of ER stress-related transcription factors bZIP60 and bZIP28, thus compromising resistance to the oomycete pathogen *Phytophthora parasitica* (*19*). The disruption of a CUE domain-containing protein also enhanced disease response in rice (*20*). OsNTL6 alleviates ER stress sensitivity and decreases disease resistance against rice blast fungus (*21*). In contrast, tunicamycin (TM)-induced ER stress increased *Arabidopsis* susceptibility to *Pseudomonas syringae* pv tomato (Pst) DC3000 (*22*). Consistent with a potential benefit to bacterial pathogens, infection with turnip mosaic virus (TuMV) induced splicing of bZIP60 mRNA in *Arabidopsis*, and the loss of function of bifunctional endoribonuclease/protein kinase IRE1 inhibits viral infection (*23*). These reports indicate that the roles of ER stress in disease responses likely differ depending on both the pathogen type and the host.

Oomycete pathogens, such as *Phytophthora sojae*, constitute a unique category of filamentous eukaryotic microorganisms that establish infection by releasing an arsenal of small secreted proteins (*24*). Amongst this repertoire of virulence determinants, the most well-studied are host cell-entering RXLR effectors (*25*). Those cytoplasmic effector proteins are not only valuable tools for identifying immune receptors (*26*), but also versatile markers for elucidating the roles of conserved signaling pathways in adaptation to biotic stress (*27*).

In this study, using the ER-targeting RXLR effector Avh131 as a molecular probe, we identified GmEBS7a, a component of the E3 ubiquitin ligase complex involved in ERAD. GmEBS7a negatively regulates plant defenses against biotic stress. The stabilization of GmEBS7a by Avh131 hyperactivates ERAD and increases ER stress sensitivity in plants. Our findings reveal a novel immune regulatory mechanism in which the *P. sojae* effector Avh131 disrupts host ERAD pathways, thereby facilitating pathogen invasion.

## Results

### Avh131 and its paralog redundantly contribute to the full virulence of *P. sojae*

Through conserved domain analysis of *P. sojae* effectors (*28*), we identified two transmembrane domain (TMD)-containing effectors, Avh131 and Avh176, which showed sequence conservation across four genotypes (Table S1). Despite varying transcript abundances across isolates, *Avh131* and *Avh176* transcripts were significantly increased (5- to 65-fold) during infection. Specifically, *Avh131* transcription is induced at 6 h post-infection, peaks at 12 h, then downregulates with a minor increase at 48 h; *Avh176* is upregulated early in infection and stabilizes after 24 h (fig. S1), indicating their critical roles in host colonization. CRISPR/Cas9-mediated knockout (KO) of *Avh131* or *Avh176* did not alter hyphal growth and pathogenicity on soybean leaves (fig. S2, A to C and Fig. 1, A and B). However, the double KO mutant Δ*Avh131/176* showed reduced virulence, as evidenced by smaller lesion sizes and decreased biomass (Fig. 1C) and unaffected hyphal growth (fig. S2, A to C). In contrast, overexpression transformants OEAvh131 (SP-mCherry-Avh131) and OEAvh176 (SP-mCherry-Avh176), which maintained normal hyphal growth (fig. S2, D to G), produced larger lesions (1.6- and 1.5-fold increases) and enhanced biomass (3.1- and 2.6-fold increases) on soybean leaves (Fig. 1D). These results suggest that Avh131 and Avh176 are virulence-essential effectors with redundant functions. Consistent with these findings, transient expression of the mature protein of two effectors (mature version) in soybean hairy roots or *Nicotiana benthamiana* leaves significantly boosted the colonization of RFP-labeled *P. sojae* or *P. capsici* (2- and 1.2-fold; Fig. 1, E and F). As a result, we selected Avh131 as a molecular probe for further studies on immune regulation.

**Fig. 1.**
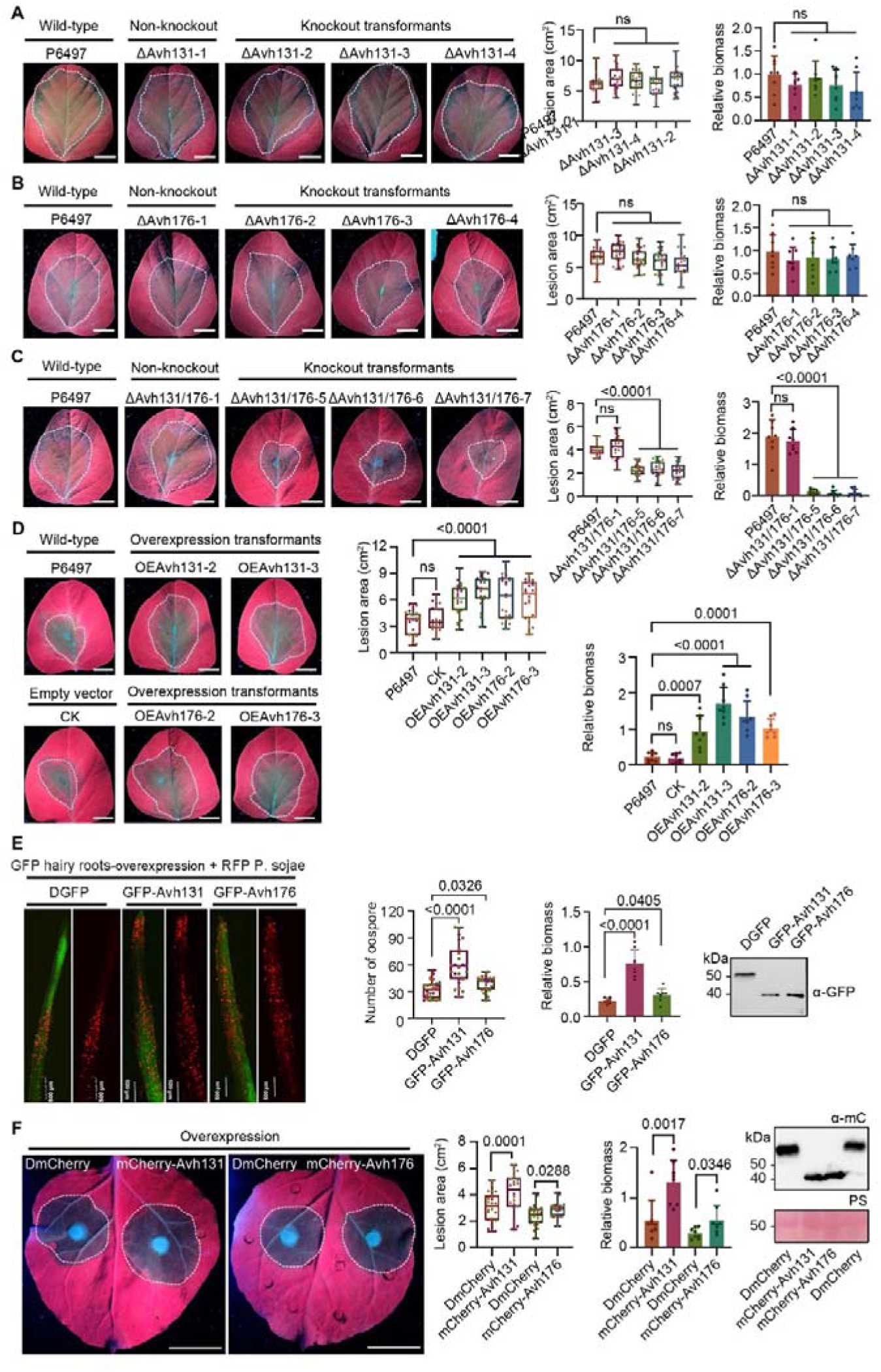
*Avh131 and Avh176* positively contribute to the virulence of *Phytophthora sojae*. **(A** to **C)** Virulence analysis of wild-type (P6497), non-knockout (Δ*Avh131*-1, Δ*Avh176*-1, Δ*Avh131*/Δ*Avh176*-1), *Avh131* knockout (Δ*Avh131*-2/3/4), *Avh176* knockout (Δ*Avh176*-2/3/4), *Avh131* and *Avh176* double knockout (Δ*Avh131*/*176*-5/6/7) transformants in leaves of soybean cultivar Williams. Scale bars, 2 cm. The lesion areas and *P. sojae* biomass were measured at 48 h post-infection (hpi). **(D)** Soybean leaves were inoculated with wild-type (P6497), empty vector (CK), *Avh131*, and *Avh176* overexpression transformants. Scale bars, 2 cm. The overexpression transformants exhibits enhanced pathogenicity, the lesion areas and *P. sojae* biomass were measured at 40 h post-infection. **(E)** Soybean hairy roots expressing double GFP (DGFP), Avh131, or Avh176 were inoculated with *P. sojae* expressing red fluorescence protein (RFP). Oospore production at 48 h post-infection (Scale bars, 0.5 mm), and *P. sojae* biomass was quantified by RT-qPCR. Total proteins of overexpression GFP[Avh131 and GFP[Avh176 in soybean hairy roots were detected by Western blotting. (**F)** *Nicotiana benthamiana* leaves transient overexpression of Avh131 and Avh176 were inoculated with *P. capsici*. Scale bars, 2 cm. The lesion areas and *P. capsici* biomass were measured at 36 h post-infection. Total proteins were detected by Western blotting. Error bars represent the mean ± standard error (In A to D, assessed with one-way ANOVA and Dunnett’s multiple comparisons, *P*<0.0001. In E and F, assessed with Student’s t test, *P*<0.05, n=24 for lesion area, n=8 for biomass). The above experiments were independently repeated three times with similar results.

### Avh131 is delivered into plant cells to enhance host susceptibility at the ER

To confirm the secretion of Avh131 in host cells, SP-mCherry-Avh131 fusion proteins were detected in both the mycelia and culture filtrate of overexpressing transformants (fig. S3A). During infection of soybean hypocotyls, SP-mCherry-Avh131 was accumulated around the haustoria-like structures of *P. sojae* transformants (fig. S3B). These findings demonstrated that Avh131 is secreted from *P. sojae*. To further demonstrate the translocation of Avh131 into plant cells, we transiently expressed vesicle marker NbAra6-GFP in *NbBAK1-*silencing *N. benthamiana* leaves, followed by inoculation with SP-mCherry-Avh131 expressing *P. sojae* transformant. Immunoblotting analysis showed that Avh131 is co-enriched with NbAra6-associated vesicles (fig. S3C), validating that a TMD-containing Avh131 is delivered into plant cells. Additionally, in fig. S3, D and E, SP-mCherry-Avh131 was exclusively detected in the apoplastic space when expressed in *N. benthamiana* leaf epidermal cells, and lost the ability to promote *P. capsici* infection. In contrast, mCherry-Avh131, which lacks the signal peptide, was localized to the intracellular space and produced larger disease lesions than the mCherry control (Fig. 1F), supporting that Avh131 must exert its function within host cells.

To characterize the subcellular localization of Avh131 in host cells and assess the functional relevance of its C-terminal TMD, co-localization assays were conducted in plant cells. The results demonstrated that Avh131 specifically accumulates at the ER, as indicated by its distinct co-localization with an ER marker (SP-mCherry-KDEL) (fig. S4A). In contrast, deletion of the TMD (Avh131^ΔTMD^) or site-directed substitution of conserved hydrophobic residues within this domain (Avh131^TMm^) resulted in diffuse cytoplasmic accumulation of the fusion proteins (fig. S4, B and C). Consistent with these subcellular localization phenotypes, both Avh131^ΔTMD^ and Avh131^TMm^ completely lost the ability to enhance plant susceptibility to *Phytophthora capsici* (fig. S4, D to F). Collectively, these findings confirm that the C-terminal TMD is indispensable for Avh131 ER anchoring and its virulence function in promoting host susceptibility.

### Soybean ER membrane protein GmEBS7a and GmEBS7b are targeted by Avh131

To dissect the mechanism of Avh131’s action in plants, we sought its interacting proteins through yeast two-hybrid (Y2H) screening. We used Avh131 as a bait to screen against a prey yeast library made from complementary DNA (cDNA) of Williams soybean infected with *P. sojae*. This screening revealed several potential interacting candidates (Table S2). We then validated these interactions through one-on-one Y2H assays (fig. S5) and evaluated the functionality of the potential targets. This process led us to identify GmEBS7a and its homologous protein, GmEBS7b, as candidate targets, both of which are ER-bound components of the ERAD Hrd1a complex. Given that Avh131, GmEBS7a, and GmEBS7b are membrane proteins, we conducted pairwise membrane Y2H assays. These assays demonstrated that Avh131 binds to GmEBS7a and GmEBS7b, while the TMD-deleted mutant Avh131ΔTMD showed a weak interaction (Fig. 2A). A co-localization analysis revealed that GmEBS7a, GmEBS7b, Avh131, and an ER marker predominantly localized to the ER (Fig. 2B).

**Fig. 2.**
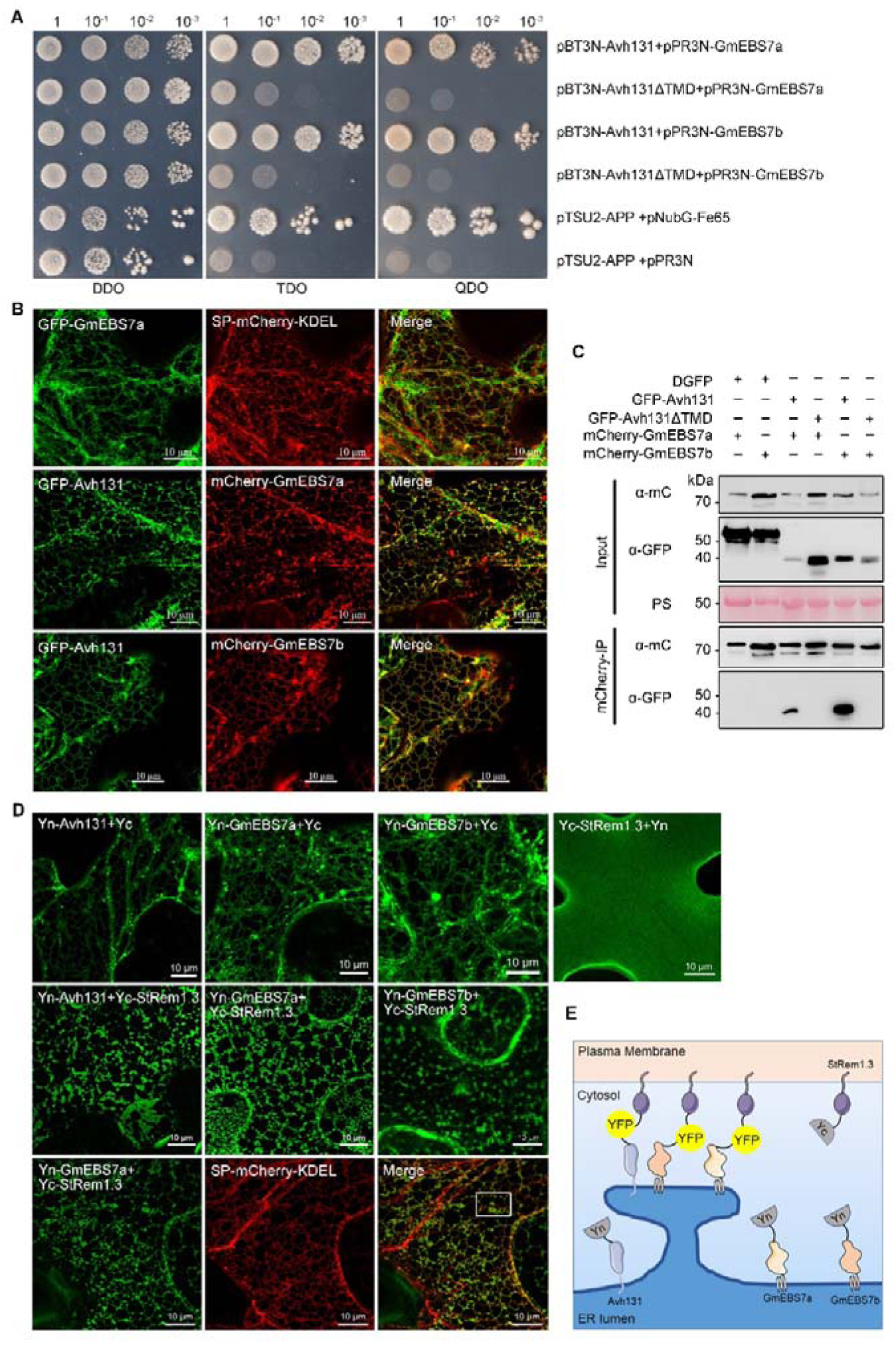
Soybean protein GmEBS7a and GmEBS7b interact with Avh131. **(A)** GmEBS7a and GmEBS7b from *Glycine max* interact with Avh131 in yeast. Yeast transformants were grown on either the non-selective medium SC/-Trp/-Leu (DDO), or on the selective media SC/-Trp/-Leu/-His (TDO) and SC/-Trp/-Leu/-His/-Ade (QDO). Ptsu2-App2 and pNubG-Fe65 were the positive control, and pTSU-App2 and pPR3-N were the negative control. **(B)** Confocal microscopy of *N. benthamiana* leaves co-expressing GFP-GmEBS7a and ER marker (SP-mCherry-KDEL), GFP-Avh131 and mCherry-GmEBS7a or mCherry-GmEBS7b. Scale bars, 10 μm. **(C)** GmEBS7a and GmEBS7b interact with Avh131 *in planta*. Proteins were extracted from *N. benthamiana* leaves expressing mCherry-GmEBS7a or mCherry-GmEBS7b with GFP[Avh131 or GFP-Avh131^ΔTMD^. Protein complexes were immunoprecipitated with mCherry beads. Co-precipitation of GFP-Avh131 was detected by Western blotting. **(D** and **E)** Bimolecular fluorescence complementation (BiFC) of Yn-Avh131, Yn-GmEBS7a, Yn-GmEBS7b and YcStRem1.3 without or with SP-mCherry-KDEL in transiently transformed *N. benthamiana* leaves. Scale bars, 10 μm. The above experiments were independently repeated three times with similar results.

To confirm the interaction *in vivo*, co-immunoprecipitation (Co-IP) was performed by expressing mCherry-GmEBS7a or mCherry-GmEBS7b with GFP-Avh131 or GFP-Avh131^ΔTMD^ in *N. benthamiana*, followed by isolation of the complex using mCherry beads. All proteins were detected in the relevant input samples. As expected, Avh131 was immunoprecipitated in the presence of GmEBS7a or GmEBS7b, but the nonfunctional mutant Avh131^ΔTMD^ was not (Fig. 2C). In addition, using a bimolecular fluorescence complementation (BiFC)-based artificial ER-PM tethering system (*29*), we demonstrated that Yn-Avh131, Yn-GmEBS7a and Yn-GmEBS7b are associated with the PM marker Yc-StRem1.3, forming either patch-like domains or a nearly continuous network after spontaneous assembly of two half fluorescent proteins (Fig. 2, D and E). This cortical ER localization pattern suggests that the N-termini of Avh131 and GmEBS7 are exposed to the cytosol. Taken together, these results support a TMD-dependent association between Avh131 and GmEBS7a, GmEBS7b at the ER.

### GmEBS7a is a susceptibility factor and is indispensable for Avh131 virulence function

Reverse transcription quantitative PCR (RT-qPCR) was used to assess GmEBS7a and GmEBS7b expression profiles after *P. sojae* inoculation. The transcript level of GmEBS7a in *P. sojae*-infected soybean leaves remained stable during the first 12 hours post-inoculation (hpi), but increased significantly at 24, 36, and 48 hpi. In contrast, GmEBS7b was not significantly upregulated during the infection stage (fig. S6). These expression patterns indicate that GmEBS7a-involved ERAD is responsive to *P. sojae* infection and are consistent with the infection-induced upregulation of Avh131.

To further clarify the roles of GmEBS7a and GmEBS7b in plant immunity, each gene was individually overexpressed, and either *GmEBS7a/b* or *GmEBS7a* was silenced in soybean hairy roots (fig. S7, A and B). The susceptibility of these transgenic lines was assessed following inoculation with RFP-labeled *P. sojae*. Soybean hairy roots expressing *GmEBS7a* produced twice as many oospores as those expressing GFP, while *GmEBS7b* expression did not affect oospore numbers. In contrast, *GmEBS7a/b* or *GmEBS7a*-silenced hairy roots exhibited significantly fewer oospores compared to the empty vector silencing control RNAi-EV (Fig. 3, A to E). Down-regulation of *GmEBS7a/b* or *GmEBS7a* in silenced lines was confirmed by RT-qPCR (fig. S7B), and the protein integrity of GmEBS7a was validated by immunoblotting (Fig. 3F). These results indicate that GmEBS7a negatively regulates soybean resistance to *P. sojae*. Supporting this conclusion, transient overexpression of GmEBS7a in *N. benthamiana* significantly increased *P. capsici* propagation, whereas GmEBS7b did not influence plant disease resistance (Fig. 3G). The approximately 70% nucleotide identity between *NbEBS7a* and *GmEBS7a* indicates that EBS7 is relatively conserved. *NbEBS7a/b* was selected for silencing in *N. benthamiana* to explore its related functions (fig. S7, C and D). Virus-induced gene silencing (VIGS) reduced *NbEBS7a/b* expression in TRV-*NbEBS7a/b* plants by 50-80% compared to TRV-*GFP* controls, without affecting plant growth (fig. S7, E and F). The size of *P. capsici* water-soaked lesions on TRV-*NbEBS7a/b* plants decreased by 40% relative to TRV-*GFP* controls (Fig. 3H). Furthermore, the overexpression of Avh131 in *NbEBS7a/b*-silencing plants failed to promote *P. capsici* infection, while Avh131 remained functional in TRV-*GFP* plants (Fig. 3, I and J). Thus, the mechanism of EBS7-associated susceptibility appears to be conserved in two solanaceous plants. Collectively, these results suggest that the virulence function of Avh131 depends on EBS7 accumulation.

**Fig. 3.**
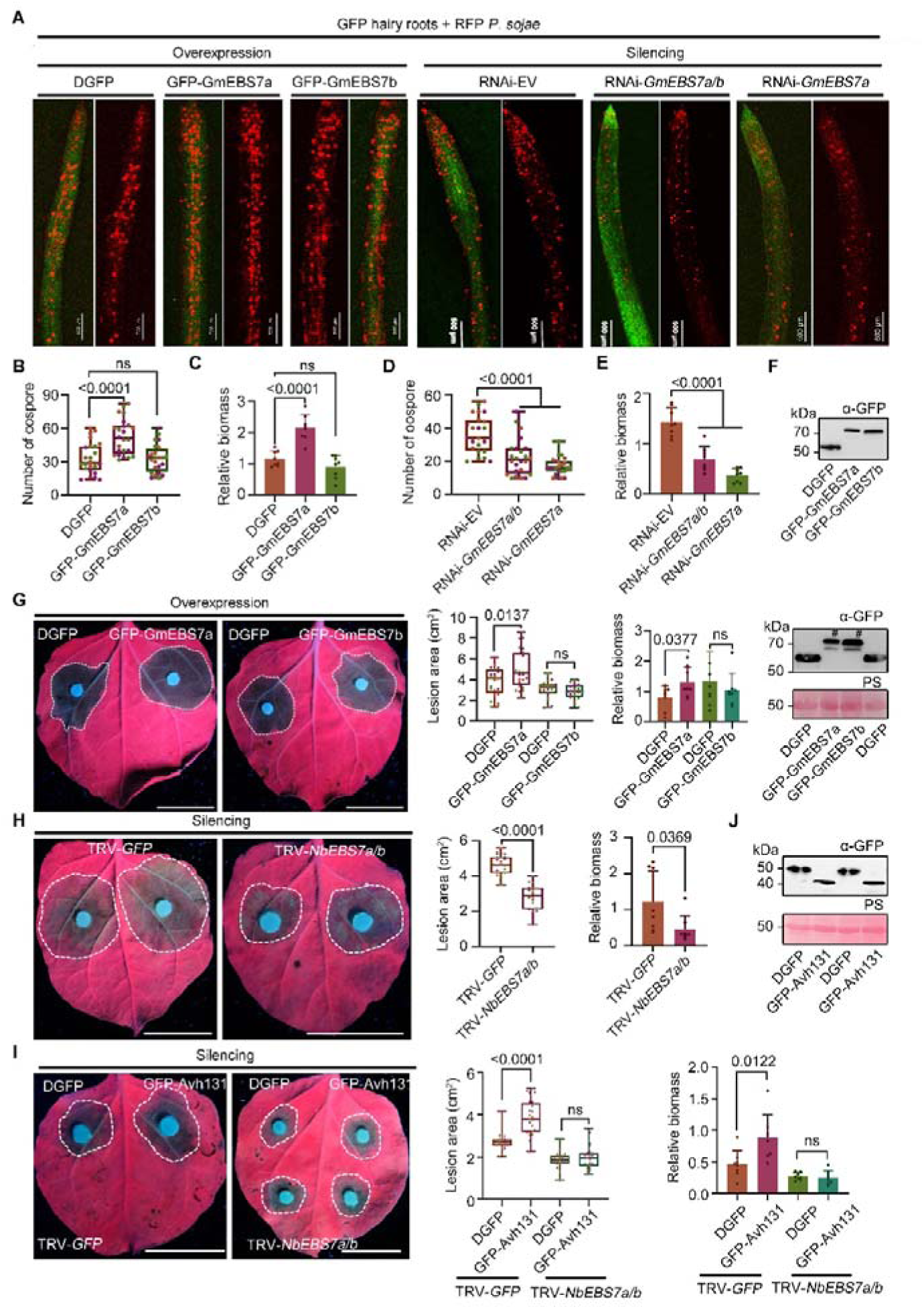
EBS7a negatively regulated plant resistance against *Phytophthora*. **(A)** Soybean hairy roots overexpressing double GFP (DGFP), GmEBS7a, GmEBS7b or silencing empty vector (EV), GmEBS7a/b, or GmEBS7a were inoculated with *P. sojae* expressing red fluorescence protein (RFP). Oospore production at 48 h post-infection (scale bars, 0.5 mm). **(B** to **E)** *P. sojae* oospore quantities (b, d) and biomass (c, e) in infected roots. *P. sojae* biomass was quantified by reverse transcription quantitative PCR (RT-qPCR). **(F)** Total protein of overexpressing DGFP, GFP[GmEBS7a or GFP-GmEBS7b in soybean hairy roots was detected by Western blotting. **(G)** *N. benthamiana* leaves transient expressionGmEBS7a, GmEBS7b were inoculated with *P. capsici*. Total protein of overexpression GFP[GmEBS7a or GmEBS7b was detected Western blotting. **(H)** *N. benthamiana* leaves silencing *NbEBS7a/b* were inoculated with *P. capsici* at 30 days post-infiltration of the silencing constructs. **(I)** *Nicotiana benthamiana* leaves with silenced NbEBS7a/b or GFP transient overexpression of Avh131were inoculated with *Phytophthora capsici*. **(J)** Total protein of overexpression GFP[Avh131 was detected by Western blotting. In B to E, assessed with one-way ANOVA and Dunnett’s multiple comparisons, *P*<0.0001, n=24 for number of oospores, n=8 for biomass. In G to I, assessed with Student’s *t* test, *P*<0.05, n=24 for lesion area, n=8 for biomass. Scale bars, 2 cm. The lesion areas were measured at 36 h post-infection. *P. capsici* biomass was quantified by reverse transcription quantitative PCR. Data and error bars were calculated from three independent experiments.

### GmEBS7a and Avh131 accelerate ERAD and trigger hypersensitivity to ER stress

To explore whether GmEBS7a or Avh131 influences ERAD, we examined the accumulation of the structurally defective but biochemically functional brassinosteroid receptor, Gmbri1-9 (*30*). The results showed that Avh131 and GmEBS7a, but not GFP controls and Avh131’s mutants, facilitate the degradation of Gmbri1-9 in *N. benthamiana* (Fig. 4A) and soybean hairy roots (Fig. 4B). A core ERAD E3 ligase GmHrd1a also enables Gmbri1-9 degradation in *N. benthamiana* (Fig. 4A). Notably, GmEBS7b does not destabilize this brassinosteroid receptor mutant (fig. S8A). A cycloheximide (CHX)-chase experiment revealed that the observed lower abundance of Gmbri1-9 was primarily due to accelerated degradation rather than decreased receptor biosynthesis, as demonstrated in Fig. 4C. Additionally, silencing *NbEBS7a*/*b* resulted in the loss of Avh131’s ability to facilitate the degradation of Gmbri1-9, as shown in Fig. 4D.

**Fig. 4.**
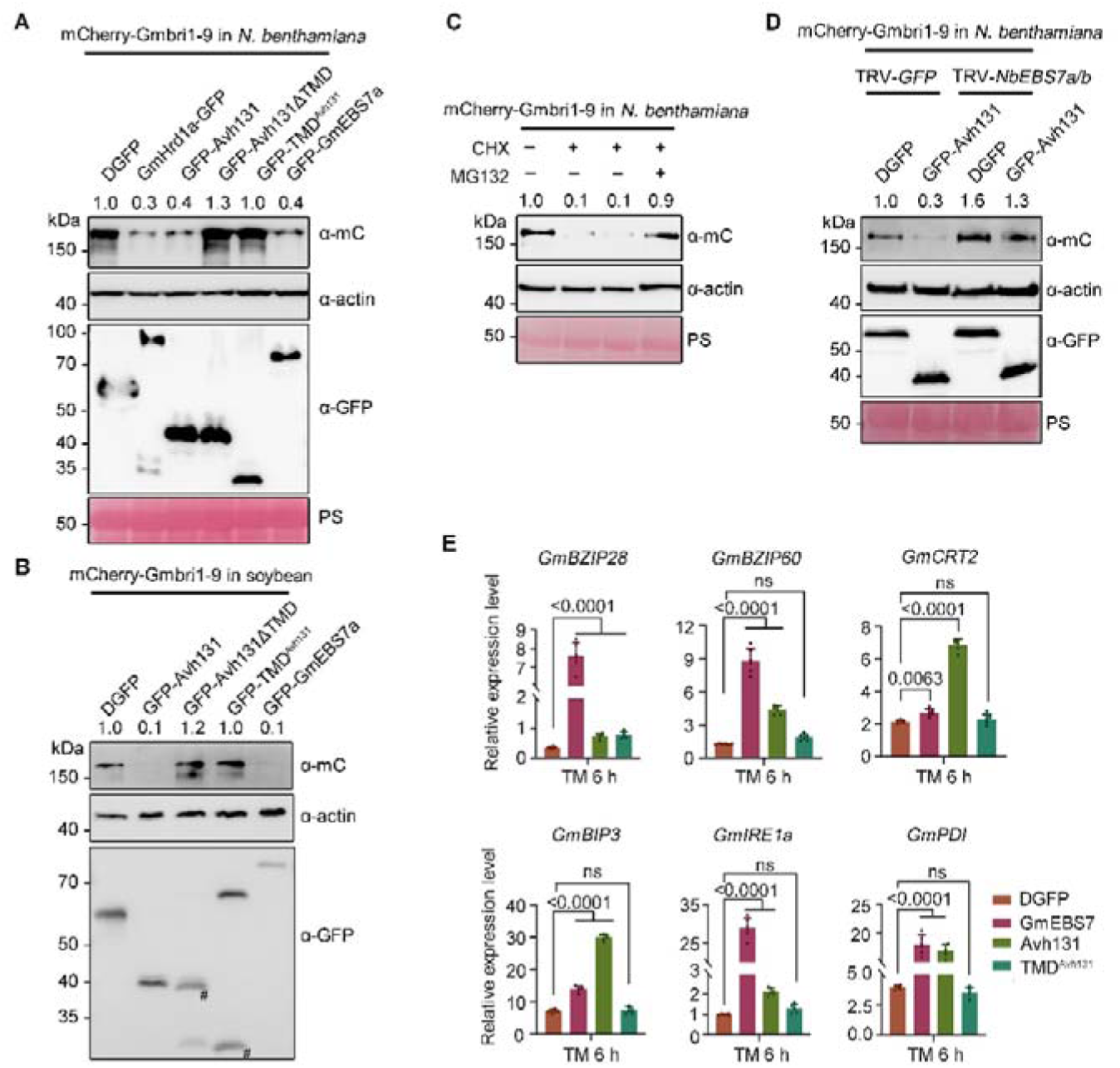
GmEBS7a and Avh131 enhance ERAD and the sensitivity to the ER stress. **(A)** Immunoblot analysis of the abundance of the protein Gmbri1-9. Total protein of transient overexpression double GFP (DGFP), Avh131, Avh131’s mutants, GmEBS7a, GmHrd1a, and Gmbri1-9 in *Nicotiana benthamiana* was detected by Western blotting. The # symbol represents the target protein band. **(B)** Total proteins extracted from transient overexpression of DGFP, Avh131, Avh131’s mutants, GmEBS7a, and Gmbri1-9 in soybean hairy roots. **(C)** Total proteins extracted from transient overexpression of Gmbri1-9 in *N. benthamiana* that were treated with 180 μM cycloheximide (CHX) or 80 μM MG132 for 12 h. **(D)** Total proteins extracted from transient overexpression of DGFP, Avh131 and Gmbri1-9 in *N. benthamiana* with silenced *NbEBS7a*/*b* or *GFP*. **(E)** The expression levels of endoplasmic reticulum stress and unfolded protein response genes were measured in soybean hairy roots overexpressing GmEBS7a, Avh131, DGFP, and TMD^Avh131^ following treatment with tunicamycin (TM: 5 μg/mL) (assessed with one-way ANOVA and Dunnett’s multiple comparisons, *P*<0.05, n=6). Total RNA was extracted from samples at 6 h post-treatment. Data shown represent fold changes of genes and display the ratio of candidate gene expression to a plant housekeeping gene. In G, the experiments were independently repeated twice with similar results. In A and D, the experiments were independently repeated three times with similar results.

To further determine whether GmEBS7a-induced susceptibility is linked to regulation of ER hematogenesis, we examined the expression profiles of several ER stress sensors and components of the unfolded protein response (UPR). In *P. sojae-*infected soybean leaves, the transcript abundance of all marker genes was significantly elevated at 24, 36, and 48 hpi (fig. S9A), indicating the ERAD pathway responds to pathogen attacks. However, agroinfiltration with Avh131 or GmEBS7a resulted in only minor changes in these markers compared to control GFP (fig. S9B), suggesting neither Avh131 nor GmEBS7a directly activates the ER stress or UPR pathways. When GmEBS7a or Avh131 was transiently expressed and followed by tunicamycin (TM) treatment, soybean hairy roots exhibited 2 to 30-fold increases in ER stress /UPR marker mRNA levels compared to the control (Fig. 4E). These data reveal that the accumulation of GmEBS7a or Avh131 leads to excessive ERAD, thereby increasing the plant’s sensitivity to TM-induced ER stress.

### Avh131 reduces GmEBS7a ubiquitination level and prevents its degradation

RT-qPCR analysis showed that Avh131 does not alter the transcription level of GmEBS7a (fig. S9C). Consequently, Avh131 is likely to affect the function or stability of the GmEBS7 protein. In line with this hypothesis, we observed an increased amount of GmEBS7a in *N. benthamiana* leaves treated with MG132 following CHX application, while the reference actin level remained stable across all treatments (Fig. 5A). Then, we examined the accumulation of GmEBS7a in *N. benthamiana* when co-expressed with Avh131, Avh131 mutants, or a DmCherry control. As shown in Fig. 5B, GmEBS7a levels were significantly elevated in the presence of Avh131 compared to co-expression with Avh131 mutants and DmCherry. MG132 treatment stabilized GFP-GmEBS7a, regardless of the co-expressed protein. Importantly, non-functional mutants Avh131ΔTMD and Avh131m did not stabilize GmEBS7a (Fig. 5 B). Similarly, Avh131 but not its mutants, stabilized GmEBS7a in soybean hairy root transgenic lines. These findings indicate that Avh131 binding is required for GmEBS7a accumulation. A substantial reduction in the ubiquitination level of GmEBS7a was also observed when co-expressed with Avh131 (Fig. 5D). On the contrary, the ubiquitination level of GmEBS7b is lower than that of GmEBS7a (fig. S8B). These results prove that Avh131 increases GmEBS7a content by reducing its ubiquitination. Furthermore, overexpression of Avh131 also extended the lifespan of the GmEBS7 interactor, GmHrd1a, and reduced its ubiquitination level (fig. S10, A and B). Notably, Avh131 did not affect the interaction between GmEBS7a and GmHrd1a (Fig. 5E), but it reduced the binding of the conjugation enzyme GmUBC32 to GmEBS7a without altering the GmUBC32 enzymatic activity (Fig. 5, F and G). These data demonstrate that Avh131 suppresses GmEBS7 ubiquitination by competitively disrupting the interaction between UBC32 and GmEBS7a.

**Fig. 5.**
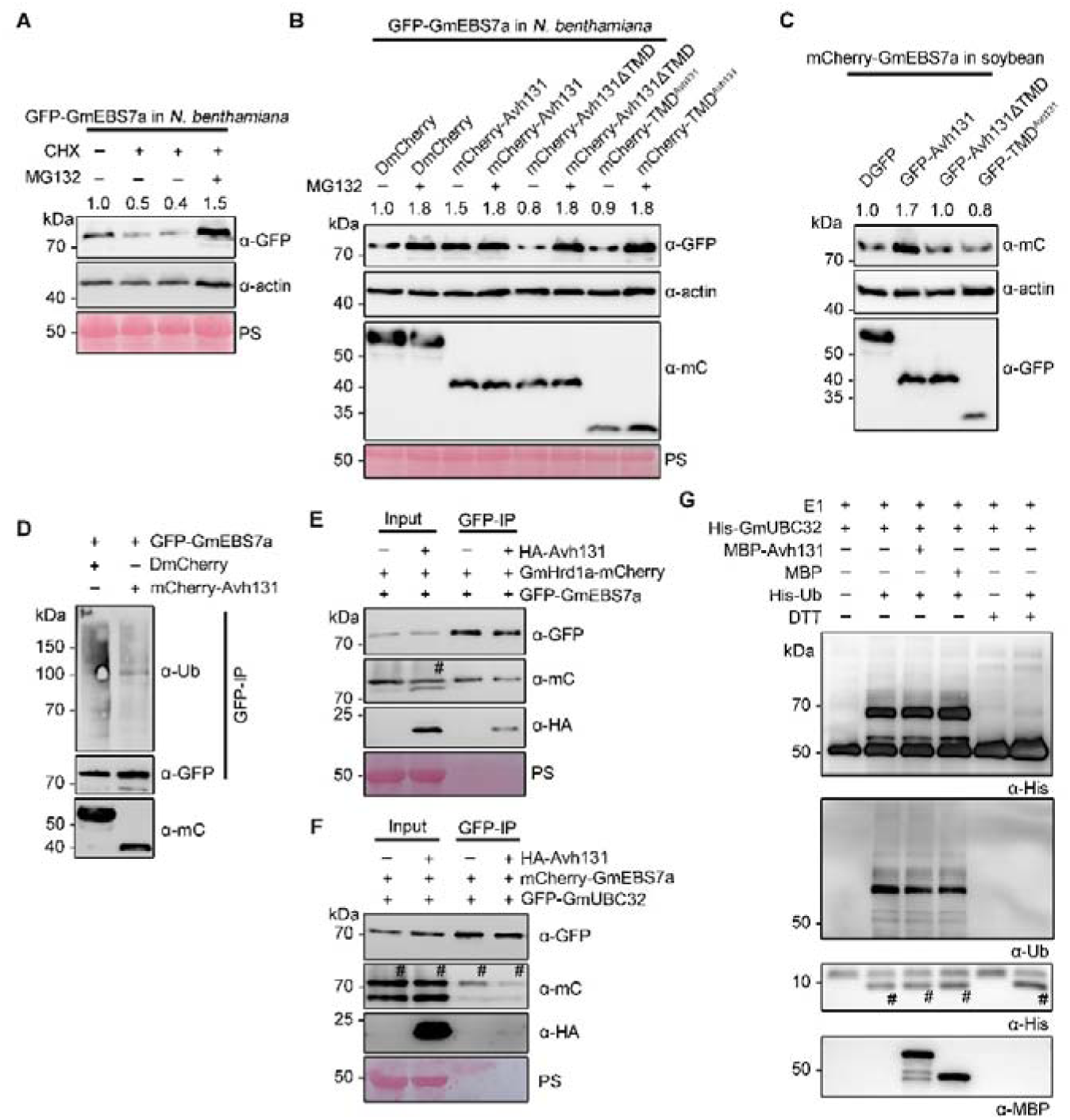
Avh131 stabilizes GmEBS7a by reducing N-terminal ubiquitination and disrupting interaction with GmEBS7a and GmUBC32. **(A** and **B)** Immunoblot analysis of the abundance of protein GmEBS7a. Total proteins were extracted from *Nicotiana benthamiana* transient overexpression of GmEBS7a, Avh131 or its mutants and subsequently treated with 180 μM cycloheximide (CHX) for 24 h or 80 μM MG132 for 12 h. **(C)** Total proteins extracted from transient overexpression of Gmbri1-9 in soybean hairy roots. In a-c, numbers on top of the immunoblot indicate relative GmEBS7a protein band intensities to actin (with the first column set to 1) as determined using ImageJ. **(D)** Immunoblot analysis of the protein ubiquitination of GmEBS7a. Transient overexpression of GmEBS7a, Avh131 in *N. benthamiana*, and subsequently treated with 80 μM MG132 for 12 h. **(E and F)** Co-IP of Avh131 with GmEBS7a and GmHrd1a or GmUBC32 in *N. benthamiana*. **(G)** E2 ubiquitin-conjugating enzyme activity of GmUBC32. The GmUBC32-Ub indicates the GmUBC32-ubiquitin adducts. DTT is a positive control confirming the specificity of thioester bond detection. In B, E, F, and G, the # symbol represents the target protein band. In D, E, and F, IP: α-GFP used to control equal amounts of proteins. The positions of molecular mass standards are indicated on the left. The PS strips show the Ponceau Red-stained RbcS band. The experiments were independently repeated three times with similar results.

Sequence analysis of GmEBS7a revealed a single predicted ubiquitination site at the N-terminal 29^th^ lysine residue. Substitution of this lysine with arginine (GmEBS7a^K29R^) markedly reduced ubiquitination (Fig. 6A). While GmEBS7a^K29R^ still accumulates at the ER, it does not promote infection when overexpressed in *N. benthamiana* leaves or soybean hairy roots (Fig. 6, B to D). Moreover, GmEBS7a^K29R^ failed to promote ERAD of Gmbri1-9 in both systems (Fig. 6, E and F). A cycloheximide (CHX)-chase experiment revealed that GmEBS7a^K29R^ exhibits greater protein stability compared to GmEBS7a (Fig. 6G). These findings suggest the 29^th^ lysine is crucial for GmEBS7a function, and maintaining a specific level of ubiquitination is essential for both ERAD activation and immune regulation.

**Fig. 6.**
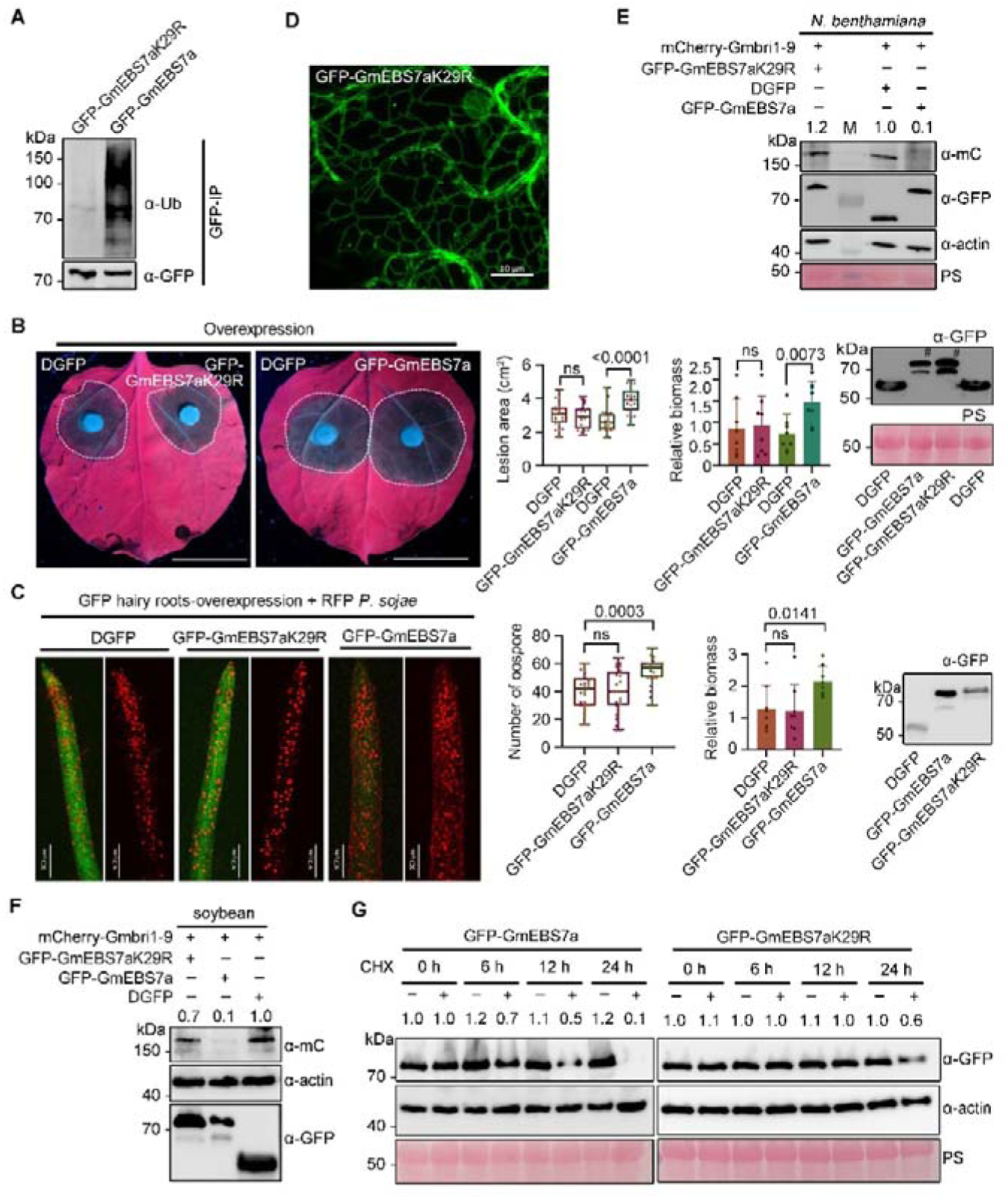
GmEBS7a^K29R^ mutant affected the ubiquitination and function of GmEBS7a. **(A)** Immunoblot analysis of the protein ubiquitination of GmEBS7a^K29R^ (the ubiquitination site mutant) in *Nicotiana benthamiana*. IP: α-GFP used to control equal amounts of proteins. **(B)** Transient overexpression of GmEBS7a^K29R^ in *N. benthamia*na did not affect *Phytophthora capsici* virulence. Scale bars, 2 cm. The lesion areas and *P. capsici* biomass were measured at 36 h post-infection. (assessed with Student’s *t* test, n=24 for lesion area, n=8 for biomass). Total protein was detected by Western blotting. **(C)** Soybean hairy roots overexpressing GmEBS7a^K29R^ were inoculated with *P. sojae* expressing red fluorescence protein (RFP). Oospore production at 48 h post-infection (scale bars, 0.5 mm). *P. sojae* biomass was quantified by RT-qPCR (assessed with one-way ANOVA and Dunnett’s multiple comparisons, n=24 for number of oospores, n=8 for biomass). Total protein was detected by Western blotting. **(D)** Confocal microscopy of *N. benthamiana* leaves overexpressing GFP-GmEBS7a^K29R^. Scale bars, 10 μm. **(E** and **F)** Effects of GmEBS7a^K29R^ on Gmbri1-9 stability in *N. benthamiana* (E) or soybean hairy roots (F). **(G)** Immunoblot analysis of the abundance of protein GmEBS7a and GmEBS7a^K29R^ from *N. benthamiana* transient overexpression of GmEBS7a, GmEBS7a^K29R^ and subsequently treated with 180 μM cycloheximide (CHX) for 0, 6, 12, 24 h. In E, F, and G, numbers on top of the immunoblot indicate relative Gmbri1-9, GmEBS7a protein band intensities to actin (with the first or DGFP column set to 1) as determined using ImageJ. The experiments were independently repeated three times with similar results.

### Ubiquitination of GmEBS7a at the ER is required for its proteolytic cleavage

Immunoblot analysis revealed that the predicted 60 kDa GmEBS7a protein appeared as both larger (∼70 kDa) and multiple smaller bands (less than 60 kDa), likely resulting from protein modifications or proteolytic cleavage (Fig. 7A). However, treatment with endoglycosidase H (EndoH) and peptide-N-glycosidase F (PNGase) did not produce a noticeable reduction in protein size (fig. S11A). Detection of the same GmEBS7a isoform following transient expression in *N. benthamiana* in the presence of the N-glycosylation inhibitor TM further supports this observation (fig. S11B). These suggest that the larger isoform of GmEBS7a is not attributable to glycosylation. As shown in Fig. 7A, co-expression with Avh131 significantly decreases the quantity of a ∼50 kDa GmEBS7a intermediate, suggesting the inhibition of specific protein cleavage. Subsequent immunoprecipitation mass spectrometry (IP-MS) analysis revealed that proteolysis occurs at the 140^th^ and 141^st^ amino acids (Table S3), suggesting cleavage on the cytosolic side near the first TMDs of GmEBS7a (Fig. 7B); This implies the involvement of site-1 protease (S1P) and site-2 protease (S2P) in GmEBS7a cleavage. Mutagenesis experiments confirmed that S1P and S2P are responsible for intermediate processing (Fig. 7C). S1P or S2P cleavage site mutants, and silencing *NbS1P* or *NbS2P*, both decrease the amount of a ∼50 kDa GmEBS7a intermediate (Fig. 7, D and E). The silencing efficiency of *NbS1P* or *NbS2P* in the silenced lines was verified by RT-qPCR (fig. S12, A to D).

**Fig. 7.**
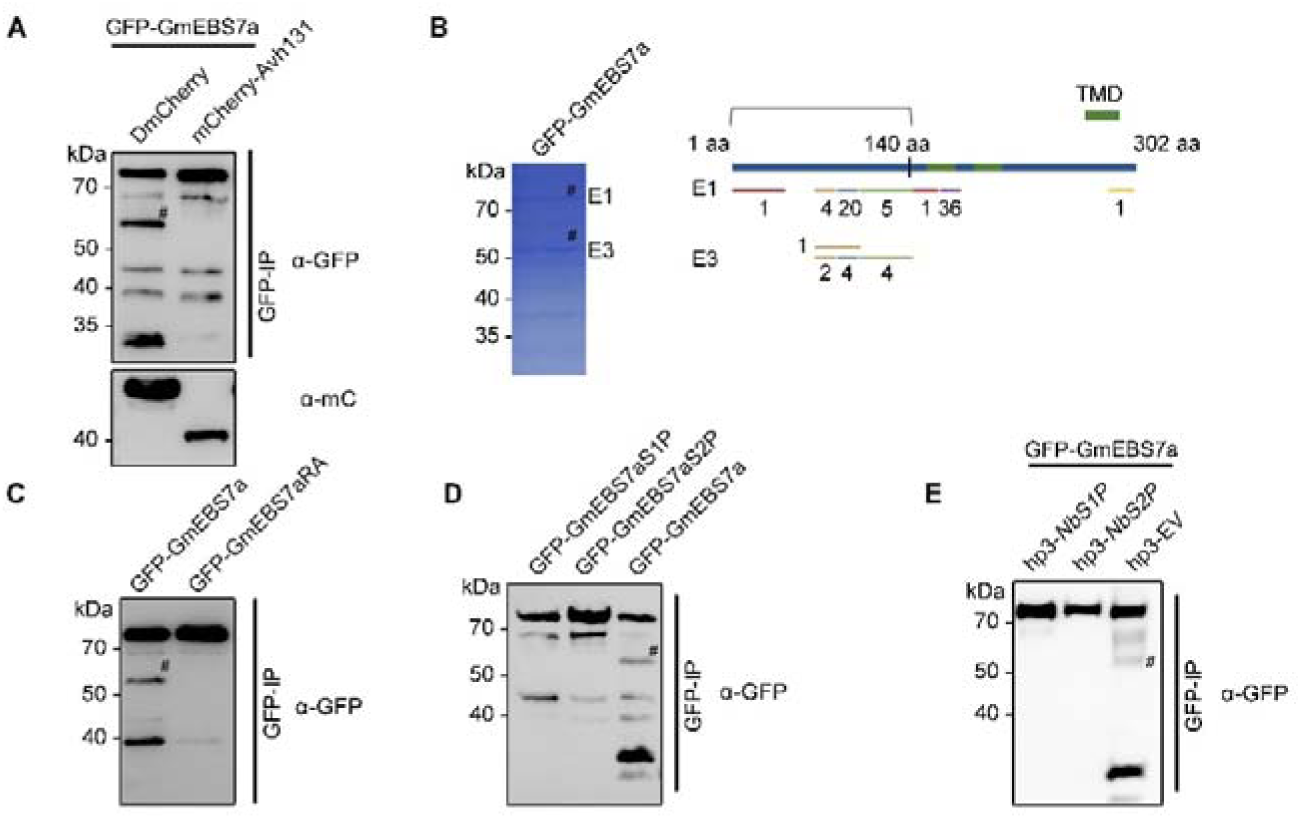
Proteolytic cleavage of GmEBS7a was reduced by Avh131. **(A)** Detected GmEBS7a protein bands from *Nicotiana benthamiana* transient overexpression of GmEBS7a and Avh131. **(B)** GmEBS7a protein bands were detected using immunoprecipitation-mass spectrometry (IP-MS) (E1: ∼70 kDa, E3: ∼50 kDa). Integrated PSMs and Sequest HT scores, selecting peptides with PSMs >2 and Sequest HT >2 for comparative analysis. The short lines represent the position and number of peptide segments in IP-MS. **(C)** Detected GmEBS7a protein bands from transient overexpression of GmEBS7a or GmEBS7a^RA^ in *N. benthamiana*. GmEBS7a^RA^ is a multiple arginine-to-alanine mutant, with mutation sites including R140/R145/R146/R174/R176/R214/R221A. **(D)** Detected GmEBS7a protein bands from transient overexpression of GmEBS7a, GmEBS7a^S1P^ (R174/R176A), and GmEBS7a^S2P^(R140/R145/R146/R214/R221A) in *N. benthamiana*. **(E)** Protein bands of GmEBS7a were detected in *NbS1P*-, *NbS2P*-, or empty vector (EV)-silenced *N. benthamiana* leaves following transient overexpression of GmEBS7a. In A-E, the # symbol represents the target protein band. The positions of molecular mass standards are indicated on the left. The PS strips show the Ponceau Red-stained RbcS band. The experiments were independently repeated three times with similar results.

Interestingly, we found that the S1P-S2P-resistant mutant GmEBS7a^RA^ resides in the ER (Fig. 8A) and retains the capacity to activate ERAD and enhance plant susceptibility (Fig. 8, B to F). In contrast, mutant GmEBS7a^K29R^ is also found in the ER but lacks these activities. Mutation of the 29^th^ lysine residue completely abolishes proteolytic cleavage (Fig. 8G) and generates a single ∼70 kDa GmEBS7a band from both total and microsome protein (Fig. 8H). This confirms that the ubiquitination site at the N-terminus determines GmEBS7a hydrolysis at the ER. In contrast, cleavage-resistant GmEBS7a^RA^ could be ubiquitinated (fig. S13), suggesting the ubiquitination at the cytosolic terminus is a prerequisite for hydrolysis, ER-releasing, and proteasome-mediated degradation.

**Fig. 8.**
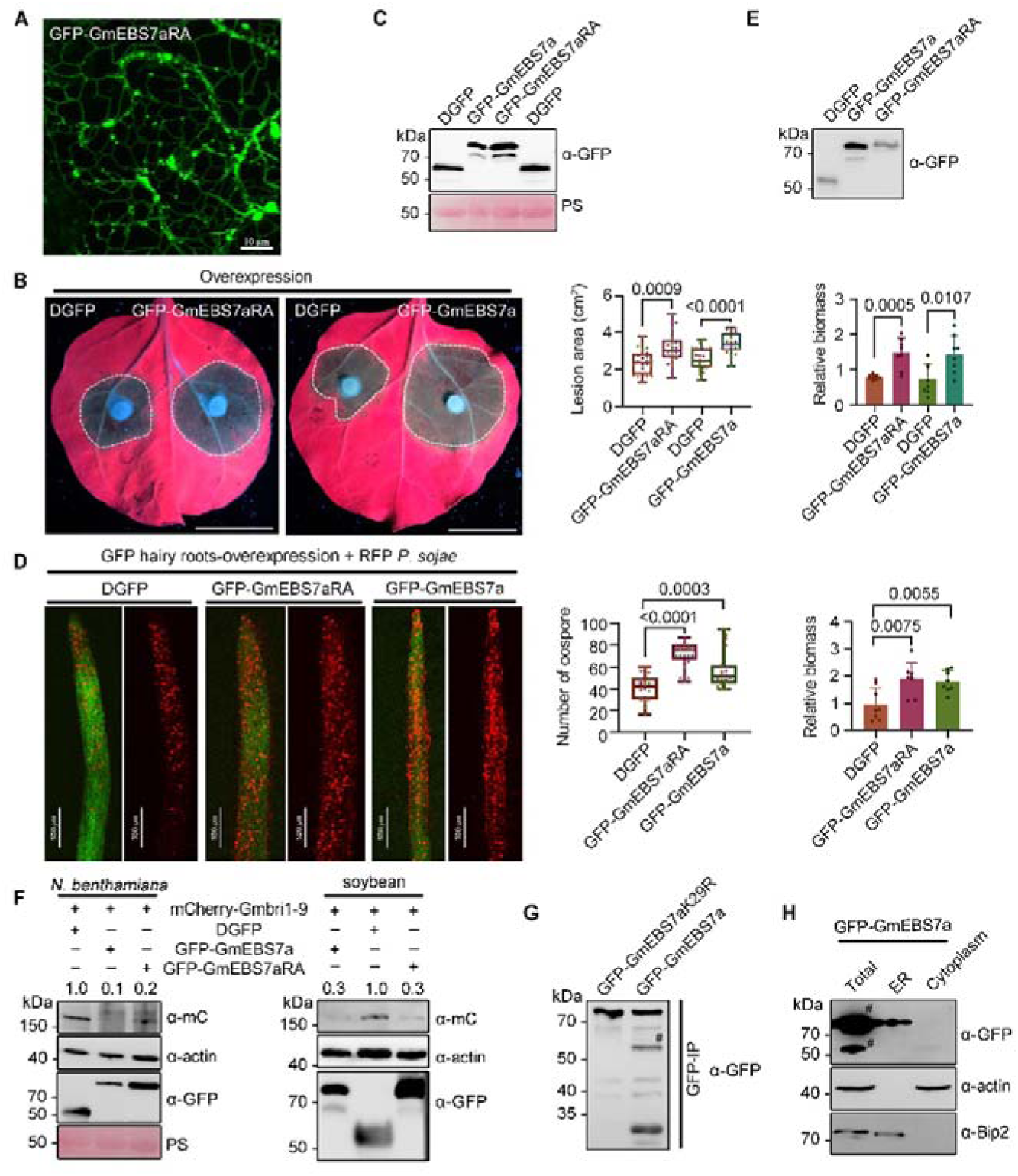
GmEBS7a^RA^ mutant affected proteolytic cleavage of GmEBS7a but unaffected the function of GmEBS7a. **(A)** Confocal microscopy of *Nicotiana benthamiana* leaves expressing GFP-GmEBS7a^RA^. Scale bars, 10 μm. **(B)** *N. benthamiana* leaves transient overexpression of GmEBS7a^RA^ were inoculated with *Phytophthora capsici*. Scale bars, 2 cm. The lesion areas and *P. capsici* biomass were measured at 36 h post-infection (assessed with Student’s *t* test, *P*<0.001, n=24 for lesion area, n=8 for biomass). **(C)** Total protein of overexpression GFP-GmEBS7a^RA^ was detected by Western blotting. **(D)** Soybean hairy roots overexpressing GmEBS7a^RA^ were inoculated with *P. sojae* expressing red fluorescence protein (RFP). Oospore production at 48 h post-infection (scale bars, 0.5 mm). *P. sojae* biomass was quantified by RT-qPCR (assessed with one-way ANOVA and Dunnett’s two-tailed test, n=24 for number of oospores, n=8 for biomass). **(E)** Total protein of overexpressing GFP[GmEBS7a^RA^ was detected by Western blotting. **(F)** Effects of GmEBS7a^RA^ on Gmbri1-9 stability in *N. benthamiana* (left side) or soybean hairy roots (right side). Numbers on top of the immunoblot indicate relative Gmbri1-9 protein band intensities to actin (with the DGFP column set to 1) as determined using ImageJ. **(G)** Detected protein bands from transient overexpression of GmEBS7a^K29R^ in *N. benthamiana*. **(H)** GmEBS7a protein bands in the endoplasmic reticulum and cytoplasm were detected in *N. benthamiana* using GFP, Bip2, and actin antibodies. The # symbol represents the target protein band. Error bars represent the mean ± standard errors. The experiments were independently repeated three times with similar results.

**Fig. 9.**
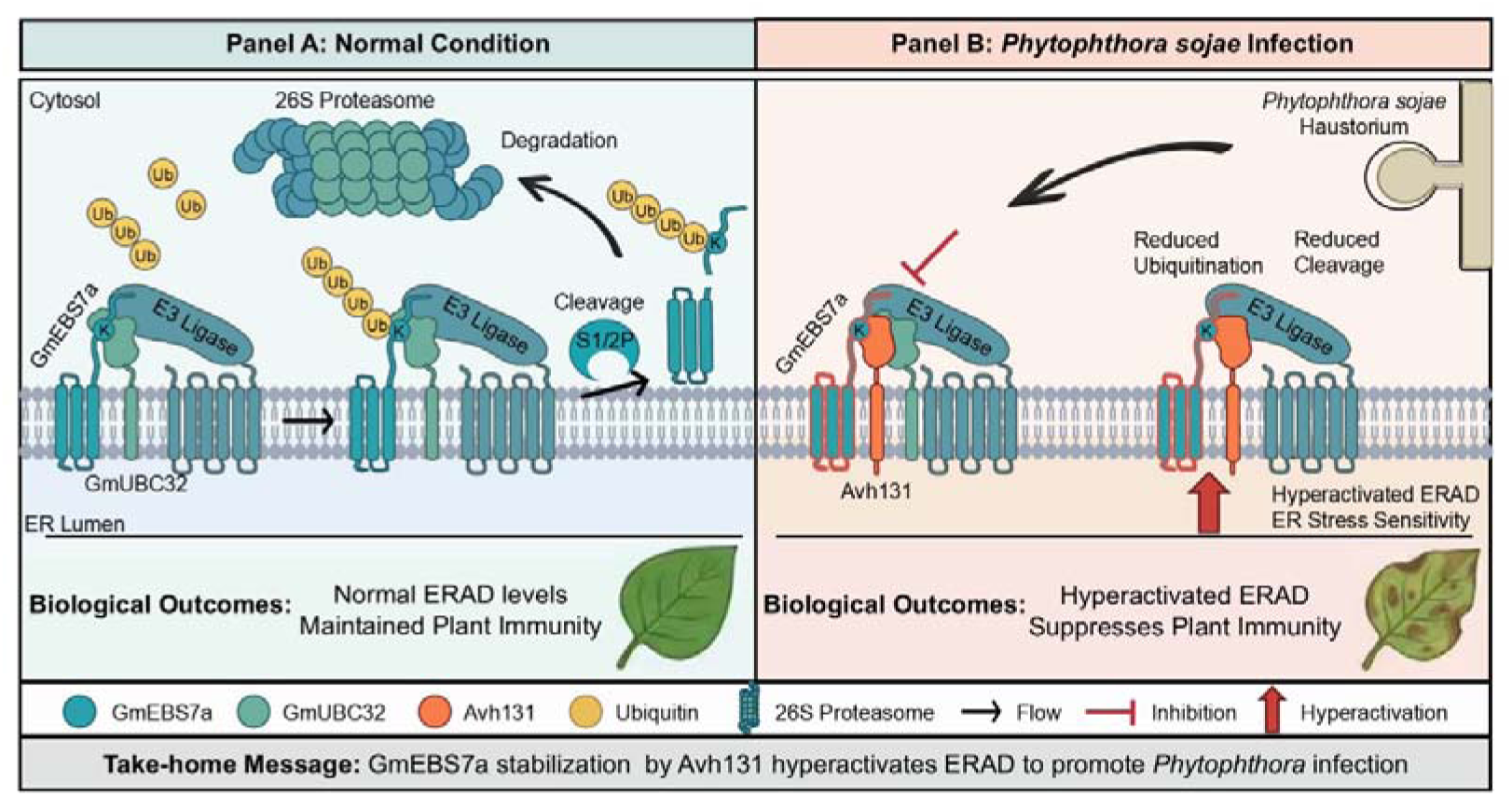
Mechanism of Avh131 mediated stabilization of GmEBS7a and Hyperactivation of ERAD suppressing plant immunity. Under normal conditions, GmEBS7a interacts with GmUBC32 (E2 ubiquitin-conjugating enzyme) and GmHrd1a (E3 ligase); GmEBS7a is ubiquitinated, cleaved, and degraded by the 26S proteasome to assist the normal functions. During infection, *P. sojae* produces haustoria, through which effectors are secreted and translocated into host cells. Avh131 associates and stabilizes endoplasmic reticulum anchored protein GmEBS7a, reducing ubiquitination and cleavage that is due to the Avh131 competitive binding of GmUBC32; at the same time, Avh131 and GmEBS7a hyperactivated the ER stress and ERAD response in the host plant. This triggers hypersensitivity to ER stress and accelerates ERAD, subsequently suppressing plant immunity.

## Discussion

This study establishes Avh131, a transmembrane domain (TMD)-containing effector from *P. sojae*, as a functional probe that targets the host endoplasmic reticulum (ER). Avh131 anchors to the ER membrane via its C-terminal TMD, physically interacts with GmEBS7a-a core component of the GmHrd1a E3 ligase complex in the ERAD pathway, and stabilizes this protein by repressing its ubiquitination. This stabilization event impairs GmEBS7a’s proteolytic cleavage and subsequent proteasome-dependent degradation. Analyses of ERAD substrate turnover dynamics and transcriptional profiling of ER stress-responsive genes collectively confirm that enhanced ERAD efficiency and heightened ER stress sensitivity synergistically facilitate *P. sojae* infection. These observations highlight ER homeostasis as a pivotal determinant of plant immunity against oomycete pathogens, and we thus propose a novel immune-suppressive mechanism: a non-classical TMD-harboring effector anchors to the ER, hyperactivates the ERAD pathway, and thereby promotes host susceptibility to *Phytophthora* species.

Secreted proteins harboring both signal peptides (SPs) and TMDs are routinely excluded from canonical effector discovery pipelines (*31*), as this domain architecture typically directs proteins to the secretory pathway or plasma membrane—a hallmark of pattern recognition receptors (PRRs) such as FLS2 (*32*). However, a suite of complementary assays corroborates that Avh131 functions as a cytoplasmic effector. Specifically, SP-mCherry-Avh131 fusion proteins accumulate around haustoria-like structures during infection and are detectable in mycelial culture filtrates, indicating that both vegetative hyphae and haustoria serve as sites for effector secretion. Transient expression of full-length Avh131-mCherry in *N. benthamiana* results in extracellular localization rather than ER targeting, suggesting that its SP retains functionality in plant cells while the TMD becomes non-functional following entry into the host secretory pathway. In BAK1-silenced *N. benthamiana* leaves inoculated with *P. sojae* transformants expressing SP-mCherry-Avh131, the effector is enriched in NbAra6-positive vesicles, a localization pattern consistent with that of RXLR effectors Pi09216 and Pi09218 (*33*), validating a conserved translocation mechanism for TMD-containing effectors. Drawing parallels to effector secretion in *Magnaporthe oryzae* and *P. infestans* (*34, 35*), we infer that Avh131 undergoes unconventional secretion independent of Golgi-mediated processing, with its SP-TMD module likely facilitating ER-to-plasma membrane transport, though the precise molecular underpinnings of this pathway remain to be elucidated.

Most characterized fungal and oomycete effectors localize to the host nucleus or cytoplasm, with only a handful of examples exhibiting organelle-specific targeting (*36, 37*). Such subcellular localization specificity is typically governed by intrinsic targeting signals, protein-protein interactions, or pathogen-induced aggregation of subcellular structures, rendering organelle-targeting effectors invaluable probes for dissecting conserved immune regulatory pathways. Our subcellular localization assays demonstrate that Avh131 anchors to the ER via its C-terminal TMD, implying that it interferes with organelle-autonomous immune signaling cascades. A precedent for this mechanism is observed with the *P. infestans* effector Pi03192, which co-localizes with its NAC transcription factor target at the ER to block nuclear translocation and suppress defense gene transcription (*38*). Notably, we confirm that Pi03192, like Avh131, is an RXLR effector equipped with a C-terminal TMD, a finding that aligns with recent reports and underscores a conserved virulence strategy among *Phytophthora* effectors (*39*).

Beyond its well-documented roles in plant development and abiotic stress tolerance (*11*), our results show that ERAD component EBS7a negatively regulates plant immunity against the hemibiotroph *P. sojae*. Avh131 fails to boost infection in *EBS7*-silenced plants, suggesting that the virulence function of Avh131 depends on EBS7. Overexpression of Avh131 or GmEBS7a does not change the expression of ER stress marker genes, indicating that the Avh131-associated pathogenic mechanism does not rely on the suppression of ER stress responses. This differs from previous reports that the effectors regulate ER immunity by either interfering with EER stress, inhibiting VPE-mediated cell necrosis, or destroying ER homeostasis(*4, 13, 15*). In stark contrast to *P. sojae* effectors such as Avh262 (which targets the ER chaperone BiP to impair protein folding) (*40*) or Avr1d (which competitively inhibits the E3 ligase GmPUB13) (*41*), Avh131 specifically disrupts the interaction between GmEBS7a and the ubiquitin-conjugating enzyme GmUBC32 (E2) without altering GmUBC32’s ubiquitin-conjugating activity, while leaving the GmEBS7a-GmHrd1a interaction intact. This unique mode of action distinguishes Avh131 from previously characterized effectors that manipulate the host ubiquitin system, highlighting the novelty of its strategy for hijacking the ERAD pathway.

Overexpression of Avh131 or GmEBS7a accelerates the ERAD-mediated degradation of Gmbri1-9, a structurally defective yet biochemically functional brassinosteroid receptor. This observation underscores a critical physiological balance governed by ERAD activity: moderate ERAD function facilitates the clearance of misfolded proteins and restoration of ER homeostasis, whereas hyperactivation drives the aberrant degradation of essential ER-resident proteins, triggering unresolved ER stress and premature programmed cell death (PCD) (*42*). This poses a critical evolutionary trade-off for host plants: suppressing ERAD to limit pathogen exploitation may compromise basal protein quality control, while maintaining constitutive ERAD activity risks enabling effector-mediated virulence. We postulate that hyperactivated ERAD promotes pathogen infection by accelerating the degradation of functionally intact PRRs, NLRs, or positive immune regulators, thereby compromising basal and effector-triggered immune responses.

Moreover, Avh131 or GmEBS7a overexpression prolongs the transcriptional induction of UPR and ER stress marker genes following tunicamycin (TM) treatment. This heightened ER stress sensitivity impedes the re-establishment of ER homeostasis and consequently dampens plant immune competence. As a global ER stress inducer, TM inhibits N-glycosylation of nascent proteins, leading to the accumulation of unfolded/misfolded proteins in the ER lumen and robust activation of the unfolded protein response (UPR)-a process characterized by the upregulation of marker genes such as BiP3, PDI, bZIP60, and IRE1 (*19, 42*). Previous studies have linked TM-induced UPR to enhanced susceptibility to *Pseudomonas syringae*, attributing this phenotype to cellular resource reallocation (prioritizing protein folding over defense signaling) and potential suppression of PRR/NLR-mediated immune cascades. In contrast, Avh131 does not induce global ER protein misfolding or directly activate UPR markers (*43*). Instead, its targeted activation of ERAD represents a pathogen-specific, precision virulence strategy that attenuates immune signaling without triggering widespread UPR, fundamentally differing from TM’s non-specific ER stress induction.

Yeast two-hybrid (Y2H) and co-immunoprecipitation (Co-IP) assays confirm that Avh131 interacts with both GmEBS7a and its homolog GmEBS7b, which share 86% amino acid sequence identity yet exhibit distinct functional divergence in regulating plant immunity and ERAD substrate stability. A critical amino acid polymorphism, K29 in GmEBS7a versus N29 in GmEBS7b, likely underpins this functional divergence by altering post-translational modification (PTM) profiles and downstream signaling cascades. The EBS7 protein family is evolutionarily conserved across land plants: Arabidopsis thaliana encodes a single homolog (AtEBS7), while *N. benthamiana* harbors two paralogs (NbEBS7a and NbEBS7b), both of which contain predicted ubiquitination sites analogous to those in GmEBS7a. Previous studies have demonstrated that AtEBS7 modulates ERAD substrate degradation and abiotic stress responses by stabilizing the E3 ubiquitin ligase Hrd1a (*11*); however, the roles of AtEBS7 and NbEBS7 in plant immunity, the biological relevance of their ubiquitination modifications, and the mechanisms by which PTMs regulate EBS7 function remain critical unresolved questions that warrant further investigation.

The ERAD pathway consists of four sequential, coupled steps: misfolded substrate recognition, retrotranslocation to the cytoplasm, ubiquitination, and proteasome-mediated degradation. Our findings demonstrate that Avh131 anchors to the ER membrane, reducing GmEBS7a ubiquitination and partially inhibiting its proteolytic cleavage. Mass spectrometry analyses further reveal that the ER-localized ubiquitination-deficient mutant GmEBS7aK29R is resistant to S1P-S2P-mediated intramembrane proteolysis and fails to stimulate ERAD activity. In contrast, the cleavage-resistant GmEBS7aRA mutant undergoes ubiquitination and induces ERAD at the ER, indicating that ER-localized ubiquitination is a prerequisite for subsequent proteolytic cleavage and proteasomal degradation. Whether endogenous GmEBS7a undergoes pathogen-induced proteolytic cleavage during *P. sojae* infection merits further investigation using GmEBS7a transgenic plant lines. The GmEBS7aK29R mutant is not only resistant to proteasome-mediated degradation but also loses its ability to elicit an ERAD-a phenotype, a phenotype that may stem from Lys29 being a key functional residue (analogous to the kinase-dead mutation in the chitin receptor OsCERK1) or from reduced ubiquitination, a prerequisite for ERAD activation and immune regulation.

In summary, our study uncovers a novel immune-suppressive mechanism in which *P. sojae* effector Avh131 anchors to the host ER via its TMD, dissociates GmEBS7a from the E2 ubiquitin-conjugating enzyme GmUBC32 to stabilize it, hyperactivates the ERAD pathway, and increases ER stress sensitivity, ultimately enhancing host susceptibility. These findings broaden our mechanistic understanding of ER-centered immune regulation during plant-oomycete interactions and open new avenues for developing innovative strategies to control *Phytophthora* diseases.

## Materials and Methods

### *Phytophthora* and plant cultivation

The *P*. *sojae* strain P6497, *P. capsici* strain BYA5, and all transformants used in this study were cultured on V8 agar plates (10% V8 juice with 0.01% w/v CaCO_3_) at 25°C in the dark. *N. benthamiana* plants were grown in a glasshouse at 22°C under a 14 h/10 h day/night photoperiod at 60% relative humidity, and ∼200 μmol^−2^ s^−1^ light intensity for 4-6 weeks. Soybean plants were grown in a glasshouse at 28°C under a 16 h/8 h day/night photoperiod and a light intensity of ∼300 μmol m^−2^ s^−1^ for 8-10 days.

### Sequence Analysis

All the sequences of Avh131 and Avh176 were obtained from the FungiDB database of *P. sojae*. The full-length sequences in Supplementary Table 1. The GmEBS7a and GmEBS7b sequences were obtained from the NCBI database of *Glycine max*. The sequences of GmEBS7a and GmEBS7b were analyzed for glycosylation sites using Protter and the NetNGlyc 1.0 Server. Analyzed for ubiquitination sites using ProP 1.0-DTU Health Tech-Bioinformatic Services and hCKSAAP_UbSite server.

### Plasmid construction

The full-length coding sequences of the *Avh131* and *GmEBS7a/b* genes were cloned using ©cDNAs from *P. sojae* and soybean cultivar Williams 82. Avh131 without a SP, Avh131 deletion mutants, and the GmEBS7a’s mutants were amplified using combinations of primers. The primers are shown in Supplementary Table 4.

For homology-directed repair (HDR)-mediated gene replacement of *Avh131* and *Avh176* in *P. sojae*, the up and down 1000 bp of *Avh131* and *Avh176* were cloned into pBS-SK L, and replaced by sequences encoding mCherry. Single guide RNAs (sgRNAs) targeting *Avh131* or *Avh176* were cloned into pYF515. For the expression of proteins in *P. sojae*, the sequences encoding the full-length (including SP) Avh131 and Avh176 were each cloned into pYF3 with a mCherry tag. For transient expression in *N. benthamiana*, the mature Avh131 and Avh176 (without SP), Avh131’s mutants, full-length GmEBS7 a, GmEBS7b, and GmEBS7a’s mutants were cloned into the expression vector pMDC-mCherry/GFP/HA respectively. For protein-protein interaction through a yeast two-hybrid system, the sequences encoding the mature Avh131 and Avh131’s mutant were each cloned into pBT3N or pGBKT7, and full-length GmEBS7a and GmEBS7b were cloned into pPR3N or pGADT7. For protein-protein interaction through bimolecular fluorescence complementation, the mature Avh131 was cloned into pBiFC-Yn, and full-length GmEBS7a and GmEBS7b were cloned into pBiFC-Yc. For expression in hairy roots, the mature Avh131 and Avh176, full-length GmEBS7a and GmEBS7b were cloned into the expression vector pMDC-GFP. For silencing by RNA interference in hairy roots, a fragment of GmEBS7a/b (bases 137-386 bp) and GmEBS7a (bases 5’UTR 1150-1242 bp) were amplified and cloned into the RNAi silencing vector pFGCA-GFP. For silencing through tobacco rattle virus (TRV)-based VIGS in *N. benthamiana*, a fragment of NbEBS7a/b (bases 269-568 bp) was amplified and cloned into pTRV2. For silencing by RNA interference in *N. benthamiana*, a fragment of NbS1P (bases 1-100 bp) and NbS2P (bases 1-100 bp) were amplified and cloned into the RNAi silencing vector pGhp3. For expression of recombinant proteins, full-length GmUBC32 and Avh131 were amplified and cloned into vectors pET22-His and pMALC2X-MBP.

### *Agrobacterium*-mediated transient expression

The *Agrobacterium tumefaciens* strain GV3101 carrying the construct was cultured for 36 h in liquid Luria-Bertani (LB) medium at 28 °C. GV3101 cells were harvested by centrifugation and resuspended in infiltration medium (10 mM MES, 10 mM MgCl_2_, and 200 μM acetosyringone) and infiltrated into 4-wk-old *N. benthamiana* leaves. For transient gene expression, *N. benthamiana* leaves were harvested for protein extraction 48 h after infiltration (OD_600_=0.1). The recombinant BiFC constructs were co-transformed into *N. benthamiana* at 48 h (OD_600_=0.5). *N. benthamiana* leaves were collected 24 h after agroinfiltration (OD_600_ = 0.1) with fresh *P. capsici* BYA5 isolate inoculation, 5 mm disks of 3 days growth medium were inoculated on *N. benthamiana* leaves.

### Confocal microscopy

A laser scanning confocal microscope LSM900 with airy scan2 (Zeiss) was utilized to precisely detect and record the fluorescent signals present within the leaves of *N. benthamiana*. For the GFP was excited at a wavelength of 488 nm with a Diode Pumped Solid State (DPSS) laser, and emissions were detected within the range of 500 and 530 nm. RFP and mCherry were excited by a 561 nm DPSS laser, and the emissions were quantified between 580 and 620 nm. Image acquisition was carried out using a C-Apochromat 40x/1.2 W objective lens. Single sections and z-stacks were collected from leaf cells and processed using the Carl Zeiss ZEN blue 3.2. Images were edited by Adobe Photoshop CS5 v. 12. The plasmolysis experiment of *N. benthamiana* epidermal cells was performed by submerging agroinfiltrated leaf disks in 30% sucrose solution for a duration of 10 minutes. Subsequently, the samples were rinsed twice with dH_2_O followed by microscopic examination.

### *P. sojae* transformation, inoculation and biomass assays

Gene replacement in *P. sojae* by the CRISPR/Cas9 system was performed using polyethylene glycol-mediated transformations (*44*). The pathogenicity phenotypes of transformant strains were detected by inoculating soybean leaves using 200 zoospores or 5 mm disks in the dark at 25°C. The inoculated leaves were photographed under UV light at 36 or 48 hpi, and the lesion area was measured using ImageJ. *P. sojae* biomass was quantified by RT-qPCR at 36 or 48 hpi. Each experiment was repeated three times, each using 10 leaves.

### Transformation of soybean hairy roots and *P. sojae* infection assays

Soybean seeds of the Williams cultivar were surface-sterilized and cultured as previously described (*45*). Soybean cotyledons were collected from 6-day-old soybean seedlings. Only unblemished cotyledons. Individual cotyledons were surface sterilized with 75% ethanol and 0.5% NaClO. A small, roughly circular (0.4 cm diameter) cut was made and transferred to an MS plate. *A. rhizogenes* K599, carrying pMDCGFP, pMDCGFP-GmEBS7a/GmEBS7b, pFGCGFP, or pFGC-GmEBS7a/GmEBS7a/b was centrifuged at 4500 rpm, resuspended in infiltration buffer (10 mM MES, 10 mM MgCl_2_, 200 μM acetosyringone, pH 5.7), and adjusted to an OD_600_ of approximately 0.4. The cell suspension was applied directly to the wounded site of the cotyledon and then incubated in a growth chamber at 25°C with a 16 h/8 h light/dark regime. The efficacy of gene silencing was examined via RT-qPCR using *GmCYP2* as the reference gene. For hairy root infection assays (*46*), 4-week-old hairy roots were infected by RFP-labeled *P. sojae* mycelia, and oospores were counted at 48 hpi.

### Yeast-two-hybrid

The pBT3N-Avh131, and pPR3N-GmEBS7a/GmEBS7b plasmids was transformed into NYM51 cells, or the pGBKT7-Avh131, and pGADT7-GmEBS7a plasmids was transformed into AH109 cells, and cultured on SD/-Leu/-Trp (DDO) medium at 30°C for 2 days. Positive colonies were then transferred to SC/-Trp/-Leu/-His (TDO), SC/-Trp/-Leu/-His/-Ade (QDO) medium at 30°C for 3 days. The colonies grew SC/-Trp/-Leu/-His (TDO), SC/-Trp/-Leu/-His/-Ade (QDO) plates containing two interacting proteins.

### VIGS assays in *N. benthamiana*

The TRV-mediated gene silencing assay was conducted as described (*47*). Briefly, pTRV constructs, including pTRV1, pTRV2-*PDS*, pTRV2-*NbEBS7a*/*b* and pTRV2-*NbBAK-1*, were transferred to *A. tumefaciens* GV3101. The mixed cell suspension was adjusted to an OD_600_ of 1.0 and was infiltrated into the two primary leaves of *N. benthamiana* at the 4-leaf stage. The gene silencing efficacy was examined via RT-qPCR using *NbEF1*α as the reference gene.

### Co-immunoprecipitation and immunoprecipitation-mass spectrometry analysis

Fusion proteins were transiently expressed in *N. benthamiana* 48 h after infiltration. Proteins were extracted using GTEN buffer (25 mM Tris-HCl pH 7.5, 1 mM EDTA, 150 mM NaCl, 10% Glycerol) with 10 mM dithiothreitol (DTT), protease inhibitor cocktail (PIC), 1 mM pregnant mare’s serum gonadotropin (PMSF), and 0.1% Nonidet P40. The leaves were frozen and ground in liquid nitrogen with 1 mL extraction buffer on ice, vortexed, and kept on ice for 30 min. They were then centrifuged at 13 000 g for 10 min at 4°C, the supernatant transferred, and centrifuged again. Next, 100 μL of the supernatant was taken and added to 100 μL 2×SDS buffer, 900 μL of the supernatant was added to 25 μL anti-GFP beads. This was kept for 1-3 h on a rotator at 4 °C, then washed 5 times with ice-cold dilution buffer, and boiled at 90°C for 10 min.

For the immunoprecipitation-mass spectrometry analyses, the bound proteins were eluted with the addition of 50 µL glycine (0.2 mol/L, pH 2.5) for 40 s and were neutralized by the addition of 5 µL Tris base (1 mol/L, pH 10.4) for neutralization. Each sample was digested in gel with sequencingLgrade modified trypsin. The lyophilized peptide fractions were re-suspended in ddH2O containing 0.1% formic acid, and 8ul aliquots of which was loaded into a nanoViper C18 (Acclaim PepMap 100, 75μm×2 cm) trap column. The online Chromatography seperation was performed on the UltiMate 3000 RSLC nano (ThermoFisher). The trapping, desalting procedure were carried out with a volumn of 20 μL 100% solvent A (0.1% formic acid). Then, an elution gradient of 5-38% solvent B (80% acetonitrile, 0.1% formic acid) in 60 min was used on an analytical column (Acclaim PepMap RSLC, 75μm×25 cm C18-2 μm 100 Å). Data was acquired using an ion spray voltage of 1.9 kV, and an interface heater temperature of 320L. For a full mass spectrometry survey scan, the target value was 3e6 and the scan ranged from 350 to 1500 m/z at a resolution of 70,000 and a maximum injection time of 100 ms. For the MS2 scan, only spectra with a charge state of 2-5 were selected for fragmentation by higher-energy collision dissociation with a normalized collision energy of 28. The MS2 spectra were acquired in the ion trap in rapid mode with an AGC target of 1e5 and a maximum injection time of 50 ms. Dynamic exclusion was set for 25 s. The MS/MS data were analyzed for protein identification and quantification using Thermo Proteome Discoverer.

### SDS-PAGE and immunobloting analysis

For stabilization assays, CHX (180 μM), MG132 (80 μM), or DMSO (a control) was individually infiltrated into plant leaves 24Lor 40 h after the agroinfiltration. Proteins were fractionated by SDS-PAGE and transferred to a 0.45 μm nitrocellulose membrane. The membrane was blocked using 5% non-fat dry milk for 1 h at room temperature at 60 rpm. Incubated with antibody at 4°C overnight, washed 3 times with PBST, once for 5 min, and then incubated with a goat anti-mouse or anti-rabbit antibody at room temperature for 1 h. The membrane was washed 3 times with PBST and then visualized by excitation at 780 and 800 nm. Antibodies used were: Anti-GFP (1:5,000; AB0005; Abways), anti-mCherry (1:5,000; AB213511; Abcam), anti-ubiquitin (1:2000; CY5520; Abways), anti-HA (1:5000; AB0025; Abways), anti-MBP (1:5000; HT701; TransGen Biotech), anti-His (1:5000; HT501; TransGen Biotech), anti-Actin (1:5000; AB2001; Abways) and anti-Plant Actin (1:5000; AC009; ABclonal). Full-size images are presented in fig. S15-17.

### Expression of recombinant proteins and E2 Activity Assays

The plasmids for protein expression were transformed into Escherichia coli BL21 (DE3). The MBP-fused or 6 × His-tagged proteins were purified using Amylose Resin High Flow (New England Biolabs, Ipswich, MA, USA) or Ni2+-charged IMAC-Sepharose 6 Fast Flow (GE Healthcare, South Plainfield, NJ, USA), respectively. The thioester assay was performed under a total reaction volume of 30 μL, with the reaction system containing 50 mM Tris-HCl, pH 7.4, 10 mM MgCl_2_, with or without 20 mM DTT, 5 mM ATP, 100 ng E1 (UBE1), 2 μg of recombinant UBC32, 2 μg of recombinant Avh131, and 10 μg His-tagged ubiquitin. The reaction was incubated at 30°C for 2 hours and terminated by SDS sample buffer boiled for 10 min. In the presence of His-tagged ubiquitin, E1, and ATP, the purified His-UBC32 protein formed an adduct with ubiquitin that was lost in the presence of the thiol-reducing agent DTT, indicating that a thioester linkage was formed between ubiquitin and UBC32.

### Quantitative RT-PCR analysis

*P. sojae* hyphae inoculating on soybean leaves were collected at 0, 3, 6, 12, 24, 36 and 48 hpi. A plant RNA kit (Omega) was used to extract total RNA. cDNA was synthesized using EasyScript® One-Step gDNA Removal and cDNA Synthesis SuperMix (TransGen Biotech). qPCR was performed in a 20 µL volume containing Taq PCR Mastermix (Tiangen), 1 µL of the resultant cDNA, and matching primers, using a CFX96 machine (Bio-Rad). Expression levels were calculated using the 2^-ΔΔCt^ method. The gene-specific primers are shown in Supplementary Table 4.

*Arabidopsis* profiling has shown that SA-triggered immunity and tunicamycin-induced ER stress upregulate genes involved in secretion, protein folding, and degradation (PDI, CRT2, BiP3, IRE1, bZIP60, bZIP28)(*22, 43*). These genes are well-used ER stress/UPR marker genes (*18, 19*). Using these six *Arabidopsis* genes as seed sequences for homology search, we identified their soybean homologs GmPDI1 (Glyma.18g273900.1), GmCRT2 (Glyma.10g147600.2), GmBiP3 (Glyma05g36600.1), GmIRE1a (Glyma.11g087200.1), GmbZIP60 (Glyma.02g161100.1), and GmbZIP28 (Glyma.03g123200.1) as ER stress markers in this study. Primer sequences for these genes are provided in Table S4.

### Glycosylation analysis of GmEBS7a

To confirm the glycosylation of GmEBS7a protein, we treated GmEBS7a protein with PNGase F or EndoH (New England Biolabs) at 37°C for 4 h, Gmbri1-9 (*11*) was used as a control. For glycosylation assays, 24 h after the agroinfiltration, the N-linked glycosylation inhibitor tunicamycin was infiltrated into plant leaves for 24 h.

### Plant plasma membrane protein isolation

Cut fresh or frozen tissue (200-300 mg) into small fragments (1-2 mm × 1-2 mm) and place them onto a centrifuge column. Use a 1 mL pipette tip to repeatedly press and compress the tissue to reduce its volume. Add 50-80 mg of tissue dissociation powder and grind the tissue thoroughly for 1-2 min. Add 100 μL of Buffer A and continue grinding. Then, fill the centrifuge column completely with Buffer A and incubate uncovered on ice for 5 min. Centrifuge at 16,000 × g for 30 s. Vortex to resuspend the pellet, and centrifuge at 700 × g for 3 min. Transfer the supernatant to a new 1.5 mL centrifuge tube and centrifuge at 4°C, 16,000 × g for 30 min (prolonged centrifugation increases yield). Collect the supernatant (containing the cytosolic fraction). Add 200 μL of Buffer B to the pellet and resuspend thoroughly by pipetting or vortexing. Centrifuge at 4°C, 7,800 × g for 5 min, and collect the pellet, which represents the organelle-enriched fraction (Invent Biotechnologies, SM-005-P).

## Data Availability and Accession Number

All data needed to evaluate the conclusions in the paper are present in the paper and/or the Supplementary Materials. The accession numbers are Avh131 (PHYSODRAFT_288873), Avh176 (PHYSODRAFT_292778)), GmEBS7a (Glyma.15g165600.1), GmEBS7b (Glyma.9g059300.1), NbEBS7a/b (Niben101Scf07827g00014.1, Niben101Scf03592g02001.1), GmHrd1a (Glyma.11g01330.1), GmUBC32 (Glyma.07g240600.1), and Gmbri1-9 (Glyma.10g207300.1).

## Acknowledgments

We appreciate Dr Yuanchao Wang (Nanjing Agriculture University) for kindly sharing RFP-labeled *P. sojae* transformants. We thank Dr Xiaoyu Qiang (Northwest A&F University) and Dr Gang Wang (Fujian Agriculture and Forestry University) for their valuable suggestions and discussion.

## Author Contributions

B.G. and X.L. designed the research; G.S., Q.L., X.Z., H.Z., S.T., and Z.L. performed the research; G.S., B.G., and Q.W. analyzed the data; B.G., G.S. H.Z. and X.L. wrote the paper; all authors commented on the article before submission.

---

**Fig. S1.**
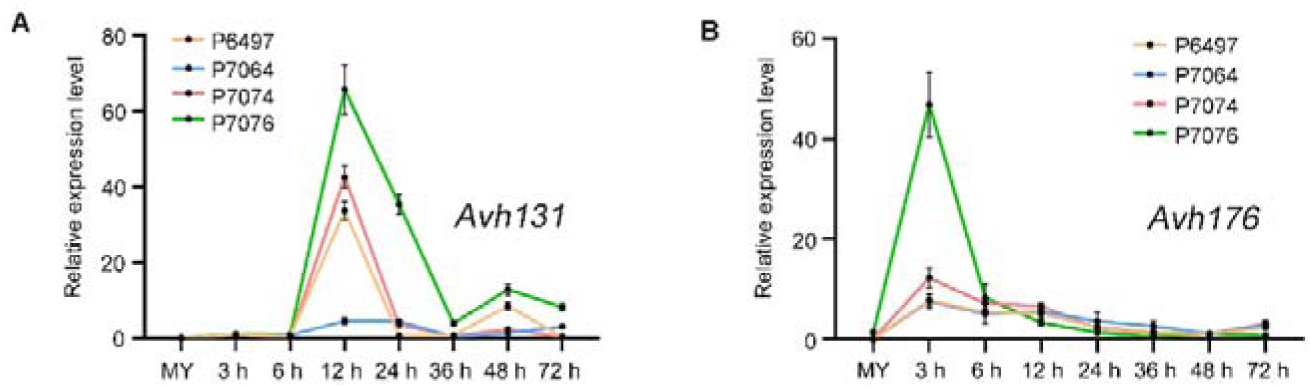
*Avh131* and *Avh176* are up-regulated during the infection. **(A** and **B)** Expression profiles of *Avh131* and *Avh176* during the infection of soybean leaves by *Phytophthora sojae* strain P6497, P7064, P7076, and P7074 were determined by reverse transcription quantitative PCR. Total RNA was extracted from mycelia (MY) or infected soybean leaves at 3, 6, 12, 24, 36, 48, and 72 h post inoculation. The *P. sojae actin* gene was used as the internal control gene. Error bars represent the mean ± standard errors. These experiments were repeated twice with similar results.

**Fig. S2.**
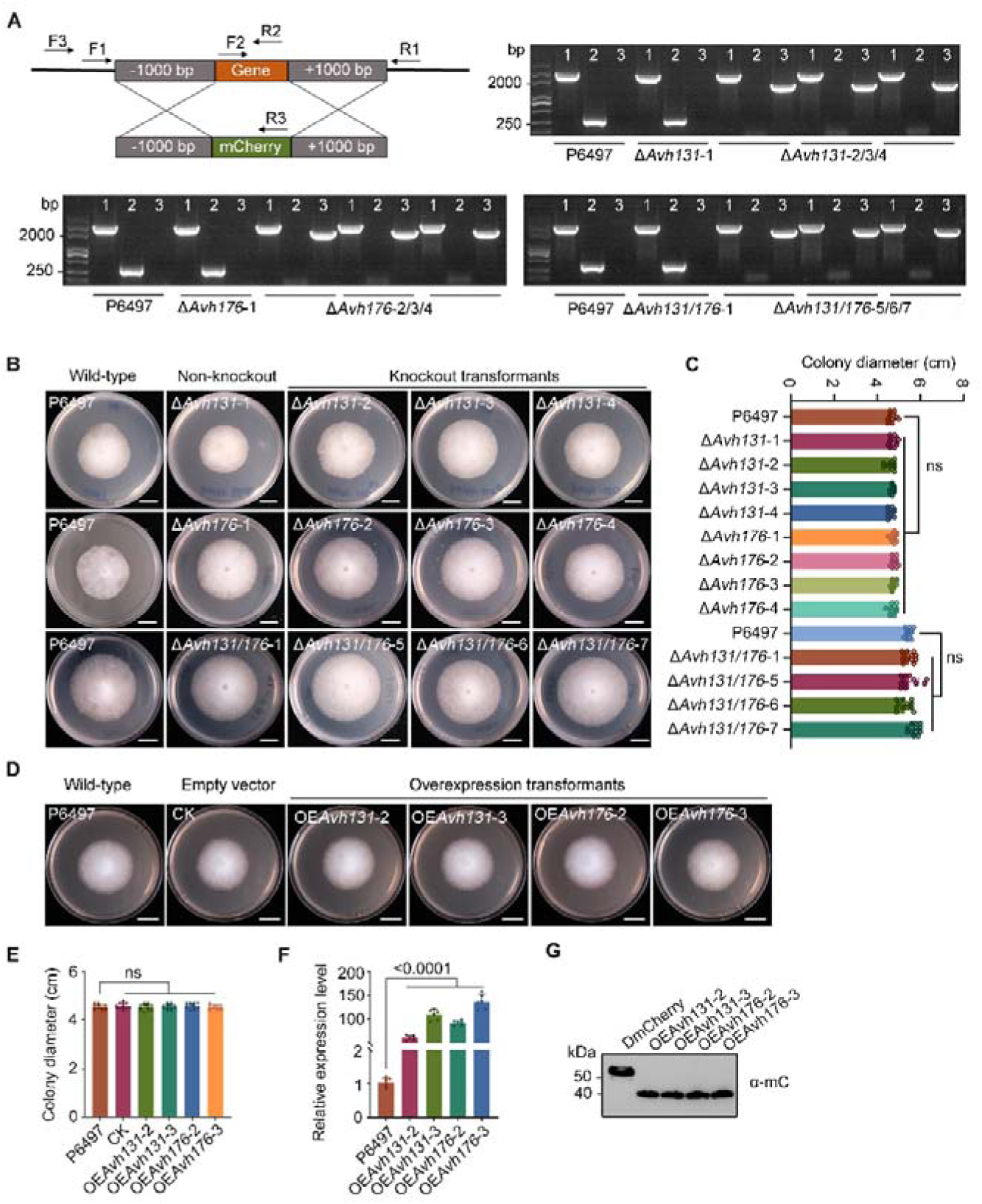
Knockout or overexpression of *Avh131 and Avh176* does not affect mycelial growth of *Phytophthora sojae*. **(A)** Model of CRISPR/ Cas9-mediated knockout of *Avh131* and *Avh176* genes knockout model. In strains Δ*Avh131*, Δ*Avh176*, and Δ*Avh131*/*176*, the relevant gene was replaced with mCherry. Primer pairs for PCR verification are marked using arrows. PCR detection of *Avh131* knockout (Δ*Avh131*), *Avh176* knockout (Δ*Avh176*), *Avh131* and *Avh176* double knockout (Δ*Avh131*/*176*) transformants. The numbers above the lanes indicate that primer pairs were used. Δ*Avh131*-1, Δ*Avh176*-1, and Δ*Avh131*/*176*-1 are non-knockout transformants. Δ*Avh131*-2/3/4, Δ*Avh176*-2/3/4, and Δ*Avh131*/*176*-5/6/7 are knockout transformants. **(B** to **E)** Wild-type (P6497), non-knockout, Avh131 knockout (Δ*Avh131*), Avh176 knockout (Δ*Avh176*), Avh131 and Avh176 double knockout (Δ*Avh131*/*176*), empty vector overexpression (CK), Avh131 overexpression (OE*Avh131*) and Avh176 overexpression (OE*Avh176*) transformants were incubated on 10 % V8 medium at 25°C for 5 days before being imaged. Scale bars, 2 cm. Colony diameters of WT and transformants were measured using ImageJ (assessed with one-way ANOVA and Dunnett’s multiple comparisons, *P*>0.05, n=12). **(F)** The transcript levels of *Avh131* and *Avh176* overexpression transformants were determined in mycelial (assessed with one-way ANOVA and Dunnett’s multiple comparisons, *P*<0.0001, n=6). **(G)** OE*Avh131* and OE*Avh176* were detected in total protein by Western blotting. Error bars represent the mean ± standard errors. These experiments were repeated twice with similar results.

**Fig. S3.**
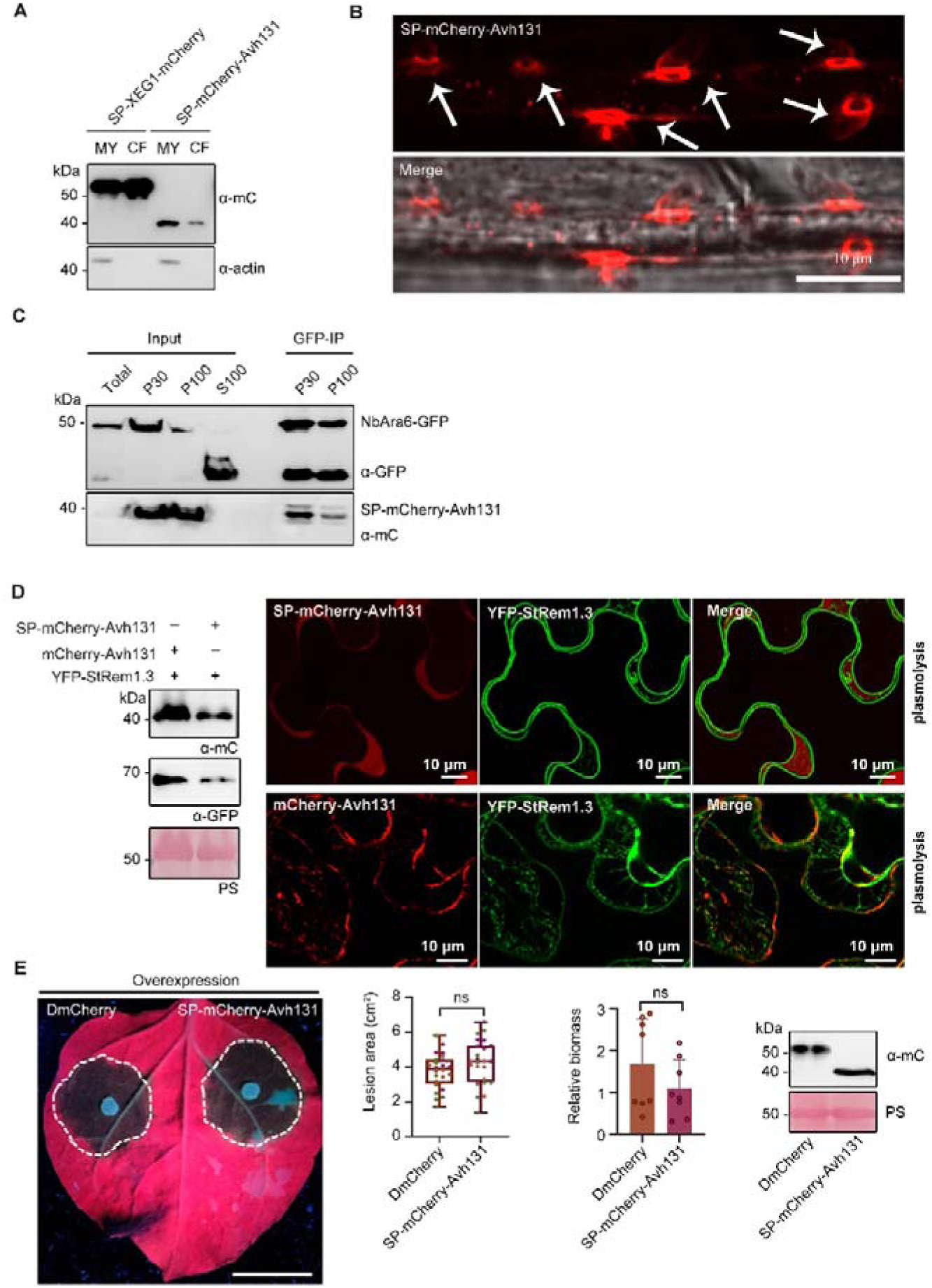
Avh131 is translocated into plant cells. **(A)** Overexpression of the Avh131 and *Phytophthora sojae* XEG1 (positive control) fusion proteins tagged with monomeric red fluorescent protein (mCherry) in the mycelium (MY) and secretion into the culture filtrate (CF) of in vitro-grown. **(B)** Localization of infection stages in etiolated soybean seedlings by *P. sojae* overexpression transformants. Scale bars, 10 μm. The arrows point to haustoria. **(C)** Plant vesicles isolated by ultracentrifugation from *NbBAK1* silencing *N. benthamiana* leaves expressing NbAra6-GFP and infected with *P. sojae* overexpression transformants SP-mCherry-Avh131. IP samples were purified NbAra6-GFP-positive vesicles in the P30 and P100 samples after incubation with GFP-Trap beads. “Total” indicates the starting material; “P30” or “P100” indicates pellets from 30,000×g or 100,000×g centrifugation. “S100” indicates supernatant from the 100,000×g centrifugation. **(D)** Transiently co-overexpressed SP-mCherry-Avh131, mCherry-Avh131 and a plasma membrane marker (YFP-StRem1.3) in *N. benthamiana* leaf cells. Leaves were then treated with 30% sucrose solution soaked for 10 minutes to induce plasmolysis, which separates the plasma membrane from the cell wall, creating a visible apoplastic gap. After plasmolysis, SP-mCherry-Avh131 was exclusively detected in the apoplastic space—distinct from the plasma membrane marker, which remained associated with the retracted cell membrane. Scale bars,10 µm. Total protein was detected by Western blotting. **(E)** *Nicotiana benthamiana* leaves transient overexpression of SP-mCherry-Avh131 was inoculated with *P. capsici*. Scale bars, 2 cm. Lesion areas and *P. capsici* biomass were measured at 36 h post-infection. Error bars represent the mean ± standard error (assessed with Student’s t test, *P*<0.05, n=24 for lesion area, n=8 for biomass). Total protein was detected by Western blotting. All experiments were repeated three times with similar results.

**Fig. S4.**
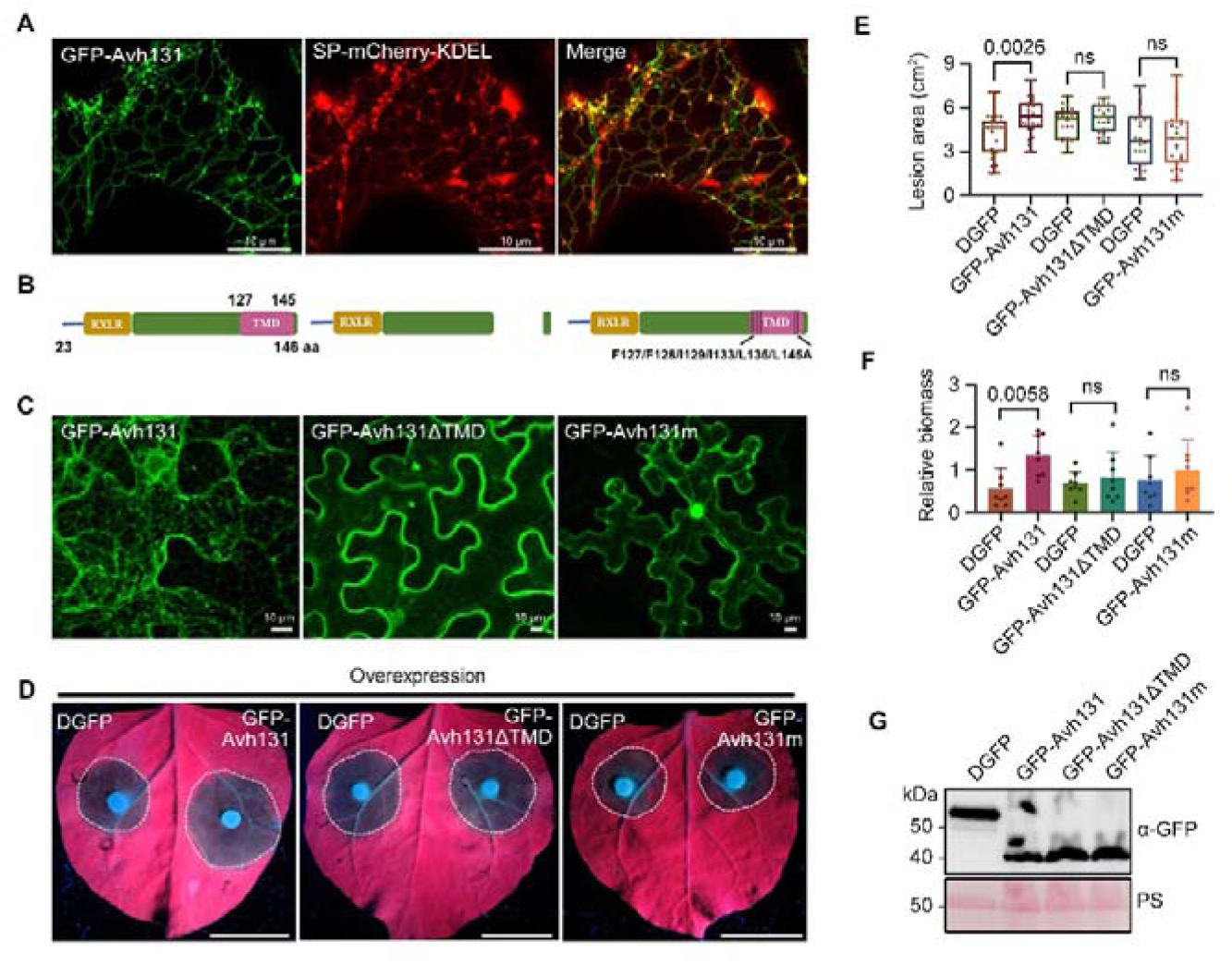
The transmembrane domain is essential for the virulence and localization of Avh131. **(A)** Localization of co-expressing GFP-Avh131 and ER marker (SP-mCherry-KDEL) by confocal microscopy at 48 h after agroinfiltration in *Nicotiana benthamiana*. Scale bars, 10 μm. **(B)** Schematic diagram showing the protein structures of Avh131 and transmembrane domain mutants. The transmembrane domain is deleted in Avh131^ΔTMD^, Avh131^TMm^ is hydrophobic amino acids in the transmembrane domain are replaced with alanine. Numbers above each construct indicate amino acid positions. **(C)** Localization of GFP-fusions of Avh131, Avh131^ΔTMD^, Avh131^TMm^ in *N. benthamiana*. Scale bars, 10 μm. **(D)** *N. benthamiana* leaves transiently expressing Avh131, Avh131^ΔTMD^, and Avh131^TMm^ were inoculated with *P. capsici*. Scale bars, 2 cm. **(E)** The lesion areas were measured at 36 h post-infection. (assessed with Student’s *t* test, *P*<0.05, n=24). **(F)** *P. capsici* biomass was quantified by reverse transcription quantitative PCR (assessed with Student’s *t* test, *P*<0.05, n=8). **(G)** Total proteins from *N. benthamiana* leaves expressing double GFP (DGFP), Avh131, Avh131^ΔTMD^, and Avh131^TMm^ were detected by Western blotting. (Assessed with one-way ANOVA and Dunnett’s multiple comparisons, *P*<0.0001, n=12). Error bars represent the mean ± standard errors. These experiments were repeated three times with similar results.

**Fig. S5.**
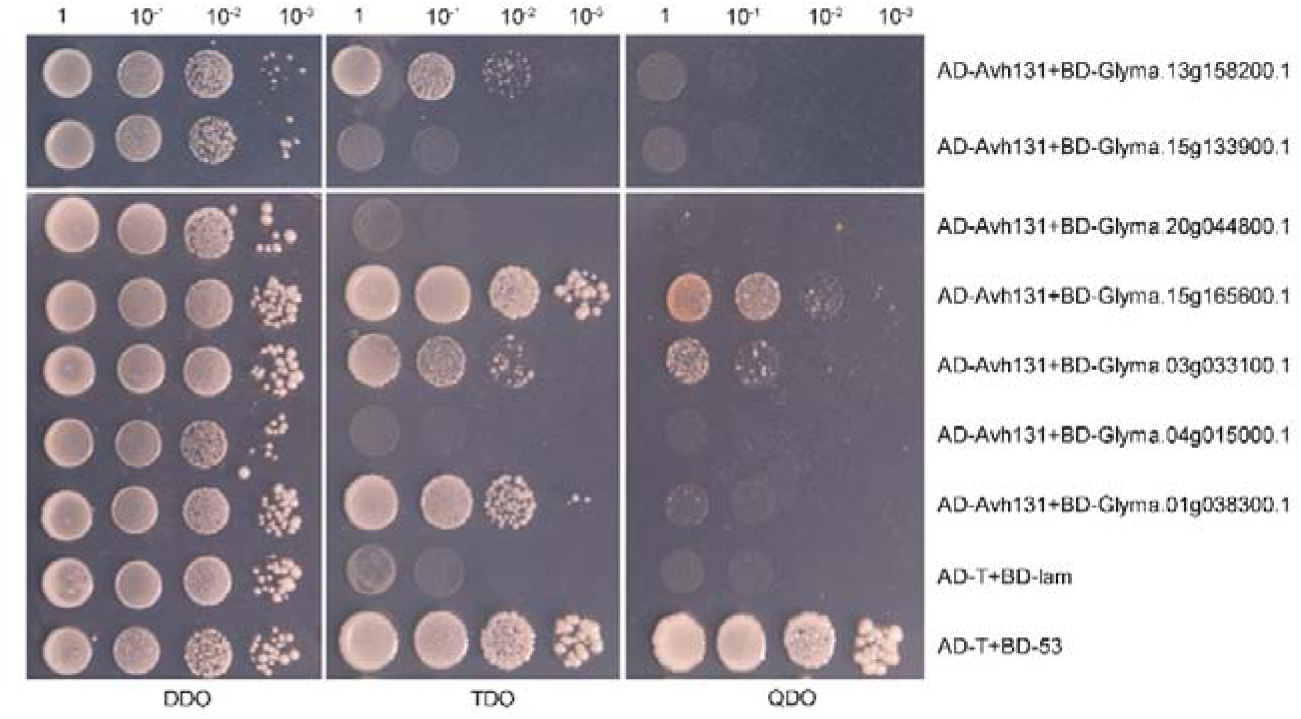
Soybean protein GmEBS7a associates with Avh131 of *Phytophthora sojae*. The one-to-one validation of interacting candidates by using nuclear yeast two-hybrid. Avh131 interacts with EBS7a of *Glycine max* in QDO. Yeast transformants were grown on the non-selective medium SC/-Trp/-Leu (DDO), or the selective SC/-Trp/-Leu/-His (TDO) and SC/-Trp/-Leu/-His/-Ade (QDO). These experiments were repeated three times with similar results.

**Fig. S6.**
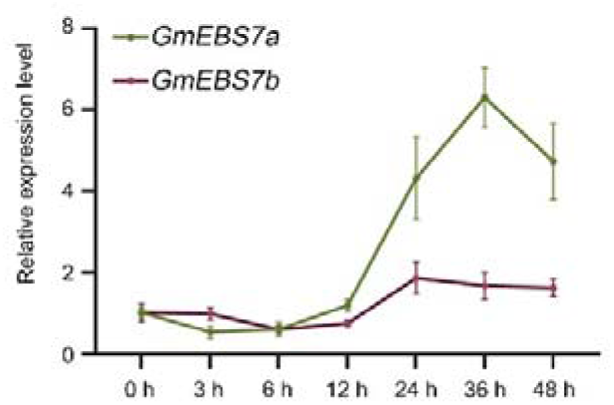
The transcript levels of *GmEBS7a* and *GmEBS7b* in soybean infected by *Phytophthora sojae*. Expression profile of *GmEBS7a* and *GmEBS7Sb* during *P. sojae* strain P6497 infection of soybean leaves. Total RNA was extracted from infected soybean leaves at 0, 3, 6, 12, 24, 36, and 48 h post-infection (with the 0 h set to 1). The *GmCYP2* gene was used as the internal control gene. The experiment was repeated twice with similar results (n[=[6, twice biologically independent samples).

**Fig. S7.**
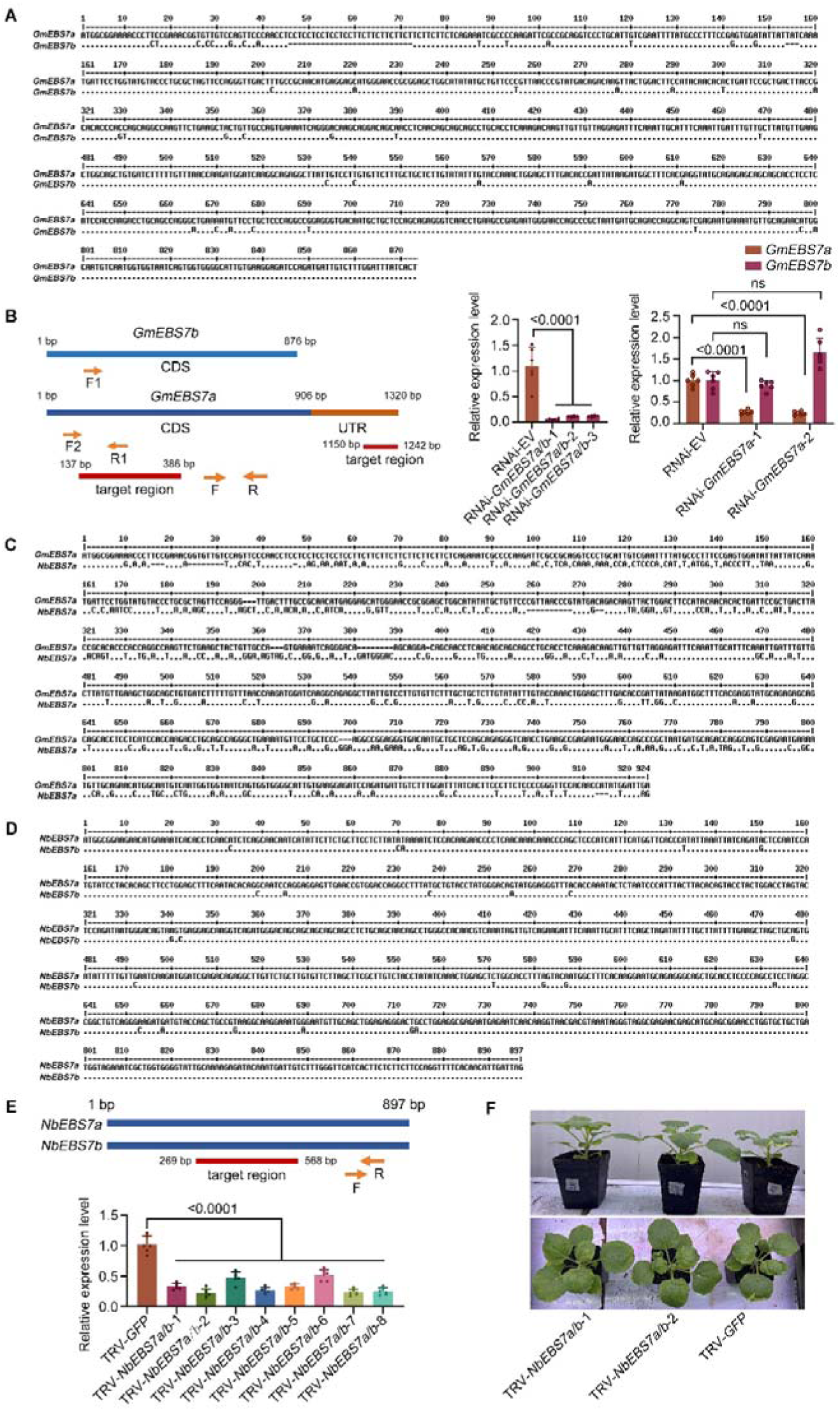
The efficiency of *EBS7* silencing in soybean and *Nicotiana benthamiana*. **(A)** Sequence alignment of *GmEBS7a* and *GmEBS7b*. **(B)** Schematic diagram of the *GmEBS7a* and *GmEBS7b* coding sequence (blue line) showing the location of the fragment (red line) used to make the hpRNAi construct and the location of PCR primers (orange arrows). The numbers represent corresponding nucleotide locations. Relative expression level of *GmEBS7a* and *GmEBS7b* in the *GmEBS7a/b* or *GmEBS7a*-silenced soybean hairy roots was determined by reverse transcription quantitative PCR (RT-qPCR) at 20-30 days post-infiltration of the silencing constructs **(C** and **D)** Sequence alignment of *GmEBS7a* and *NbEBS7a, NbEBS7a* and *NbEBS7b*. **(E)** Schematic diagram of the *NbEBS7a/b* coding sequence (blue line) showing the location of the fragment (red line) used to make the VIGS construct and the location of PCR primers (orange arrows). The numbers represent corresponding nucleotide locations. Relative expression level of *NbEBS7a/b* in *NbEBS7a/b*-silenced *N. benthamiana* was determined by RT-qPCR at 20 days post-infiltration of the silencing constructs. In B, and E, error bars represent the mean ± standard error (assessed using one-way ANOVA and Dunnett’s multiple comparisons, *P*<0.0001, n=6). The *GmCYP2* and *NbEF1*_α_ were used as the internal control gene. **(F)** Growth phenotype of *NbEBS7a/b*-silencing *N. benthamiana.* The experiments were independently repeated twice with similar results.

**Fig. S8.**
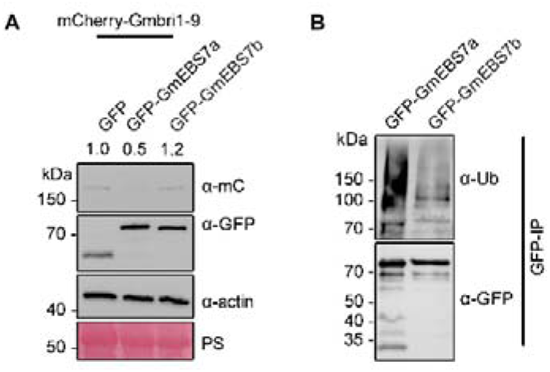
GmEBS7b lacks the functionality of GmEBS7a. **(A)** Immunoblot analysis of the abundance of the protein Gmbri1-9. *N. benthamiana* transient overexpression of GmEBS7a or GmEBS7b and Gmbri1-9, total proteins were detected by Western blotting. Numbers on top of the immunoblot indicate relative Gmbri1-9 protein band intensities to actin (with the first column set to 1) as determined using ImageJ. **(B)** Immunoblot analysis of the protein ubiquitination of GmEBS7b. IP: α-GFP used to control equal amounts of proteins. The positions of molecular mass standards are indicated on the left. The PS strips show the Ponceau Red-stained RbcS band. The experiments were independently repeated three times with similar results.

**Fig. S9.**
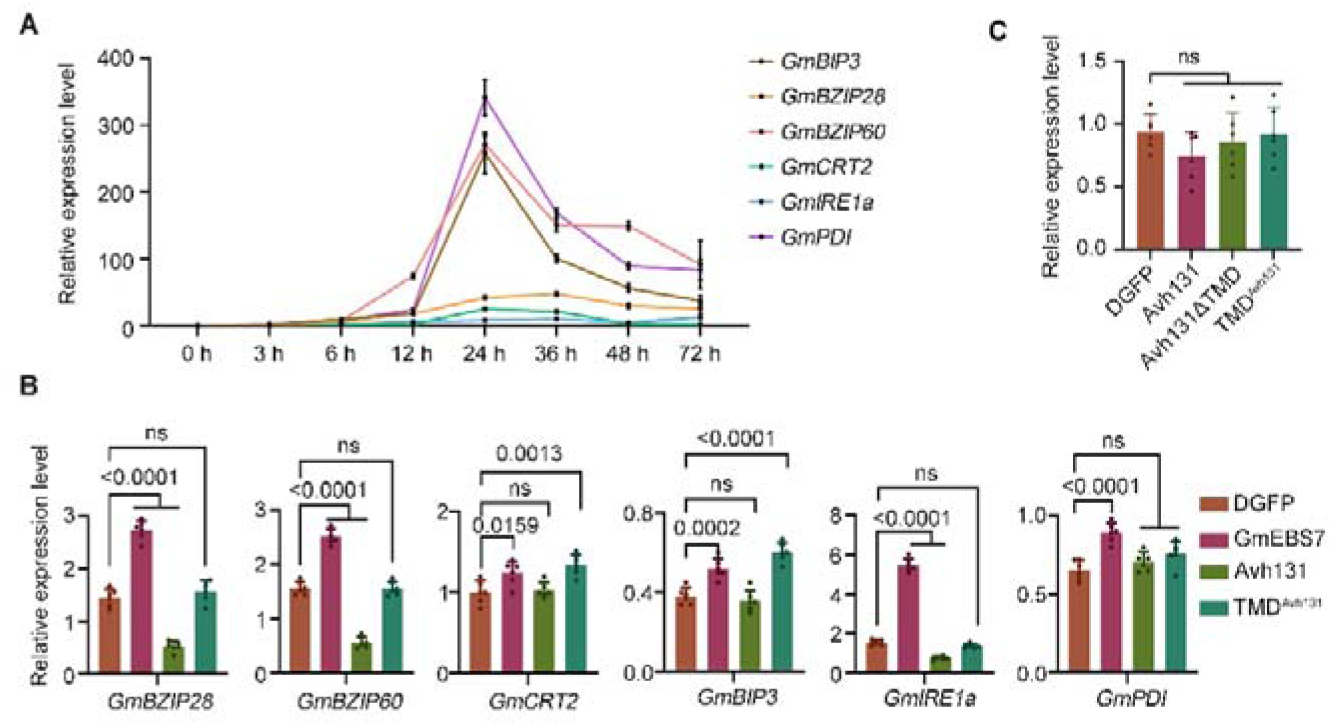
The expression level of marker genes in unfolded protein response (UPR) and endoplasmic reticulum-associated degradation (ERAD). **(A)** The expressionlevels of UPR and ERAD genes were measured by reverse transcription quantitative PCR (RT-qPCR) in soybean leaves infected by *P. sojae* strain P6497. Total RNA was extracted from infected soybean leaves at 0, 3, 6, 12, 24, 36, 48, and 72 h post-infection. **(B)** Relative expression level of UPR and ERAD genes in soybean hairy roots overexpressing double GFP (DGFP), GmEBS7a, Avh131, and TMD^Avh131^. **(C)** Relative expression level of *NbEBS7a/b* in *N. benthamia*na overexpressing DGFP, Avh131, Avh131ΔTMD, and TMD^Avh131^. Error bars represent the mean ± standard errors (assessed with one-way ANOVA and Dunnett’s multiple comparisons, *P*<0.05, n=6). The *GmCYP2* or *NbEF1*_α_ gene was used as the plant’s internal control gene. The experiments were independently repeated twice with similar results.

**Fig. S10.**
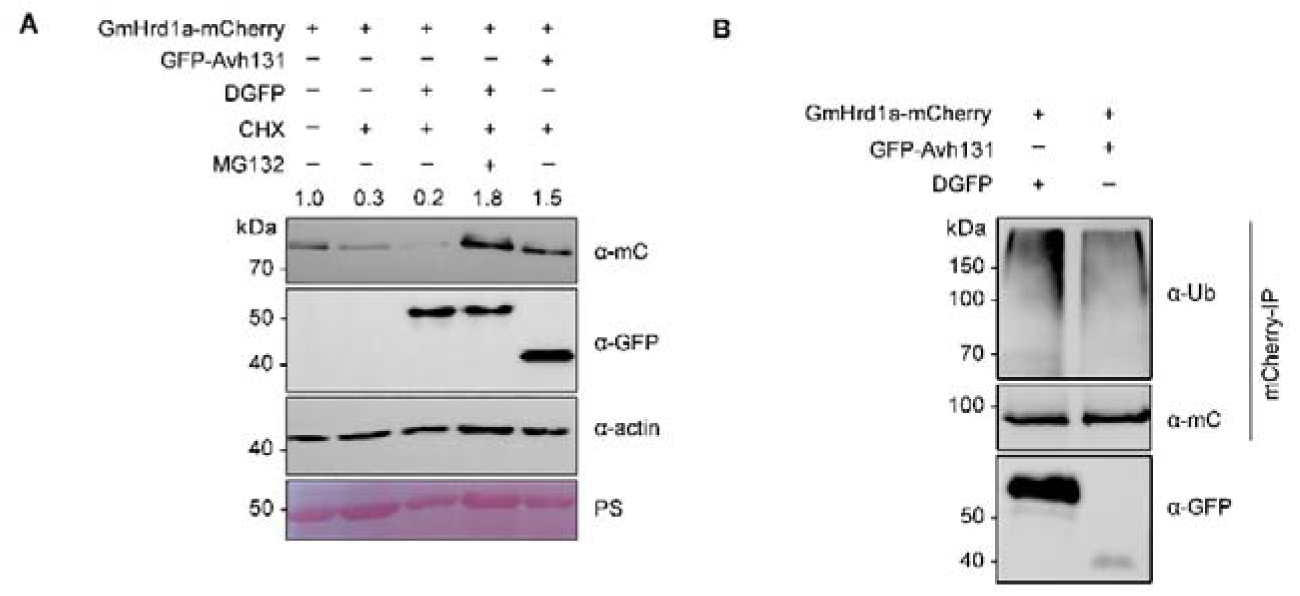
Avh131 stabilizes GmHrd1a *in planta*. **(A)** GmHrd1a was co-expressed with Avh131 or double GFP (DGFP) in *Nicotiana benthamiana* leaves and followed by 180 μM CHX or 80 μM MG132 treatment for 12 h. Total protein of GmHrd1a and Avh131 were detected by Western blotting. Numbers on top of the immunoblot indicate relative GmHrd1a protein band intensities to actin (The first column was set to 1) as determined by ImageJ. **(B)** Immunoblot analysis of the protein ubiquitination of GmHrd1a in *N. benthamiana* leaves. IP: α-mCherry used to control equal amounts of proteins. The positions of molecular mass standards are shown on the left. PS staining indicates the total protein amount. The positions of molecular mass standards are shown on the left. The experiments were independently repeated three times with similar results.

**Fig. S11.**
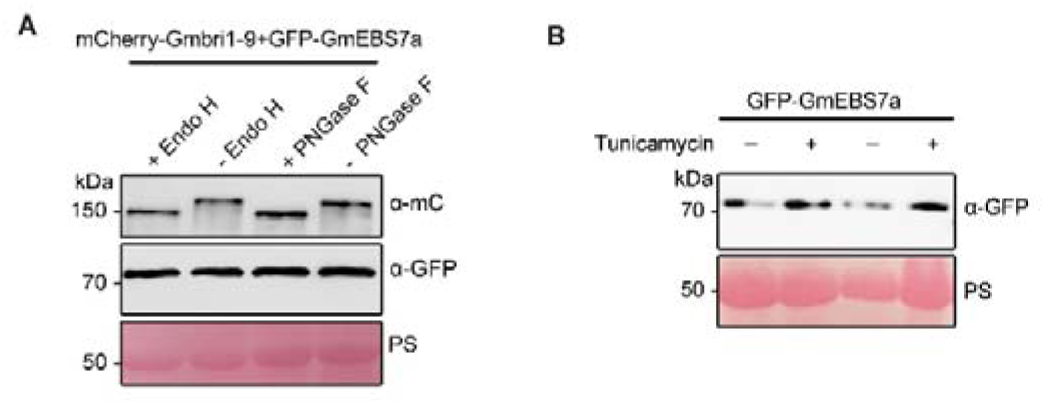
GmEBS7a did not undergo glycosylation *in vivo*. **(A)** *Nicotiana benthamiana* leaves transient overexpression of GmEBS7a and Gmbri1-9 were treated with Endo H and PNGase F for 3 h. Total proteins of GmEBS7a and Gmbri1-9 were detected by Western blotting. **(B)** *N. benthamiana* transiently expressed GmEBS7a was treated with tunicamycin (50 μg/mL) for 12 h. Protein extracts were analyzed by Western blotting using anti-GFP antibodies. PS staining indicates the total protein amount. The positions of molecular mass standards are shown on the left. The experiments were independently repeated twice with similar results.

**Fig. S12.**
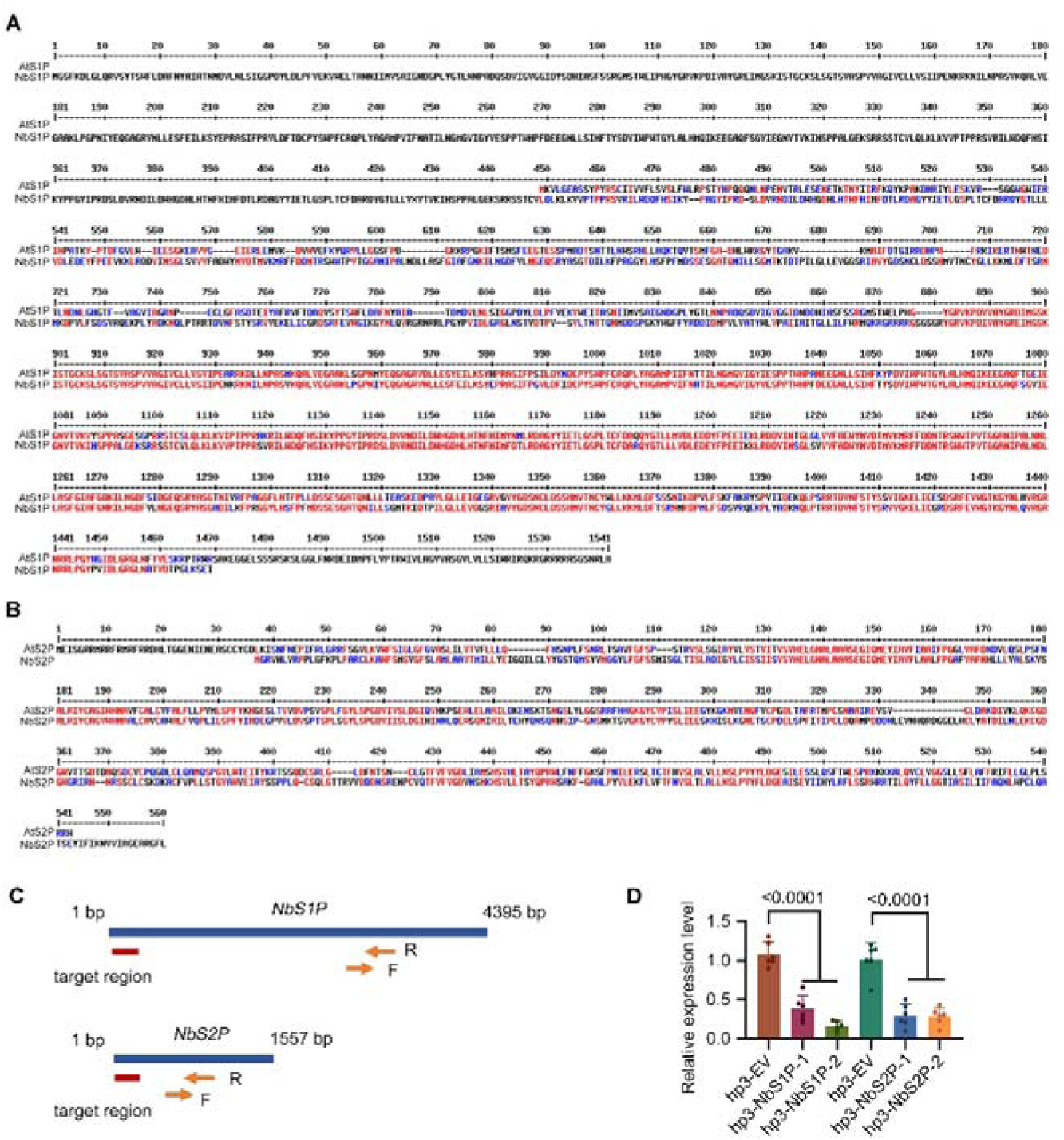
The efficiency of *NbS1P* and *NbS2P* silencing in *Nicotiana benthamiana*. **(A** and **B)** Sequence alignment of AtS1P and NbS1P, AtS2P and NbS2P. **(C)** Schematic diagram of the *NbS1P, NbS2P* coding sequence (blue line) showing the location of the fragment (red line) used to make the hpRNAi construct and the location of PCR primers (orange arrows). The numbers represent corresponding nucleotide locations. **(D)** Relative expression level of *NbS1P, NbS2P* in *NbS1P, NbS2P,* and empty vector (EV)-silenced *N. benthamiana* was determined by reverse transcription quantitative PCR at 2 days post-infiltration of the silencing constructs. The error bars represent the mean ± standard error (assessed using one-way ANOVA and Dunnett’s multiple comparisons, *P*<0.0001, n=6). The *NbEF1*_α_ were used as the internal control gene. The experiments were independently repeated twice with similar results.

**Fig. S13.**
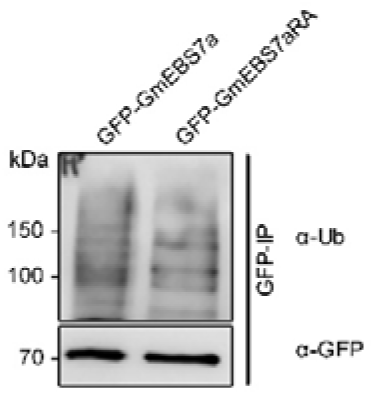
GmEBS7a^RA^ mutant did not show ubiquitination level variation. Immunoblot analysis of the protein ubiquitination of GmEBS7a^RA^ in *Nicotiana benthamiana*. α-GFP used to control equal amounts of proteins used for the ubiquitination assays. The positions of molecular mass standards are indicated on the left. The experiments were independently repeated three times with similar results.

**Table S1.**
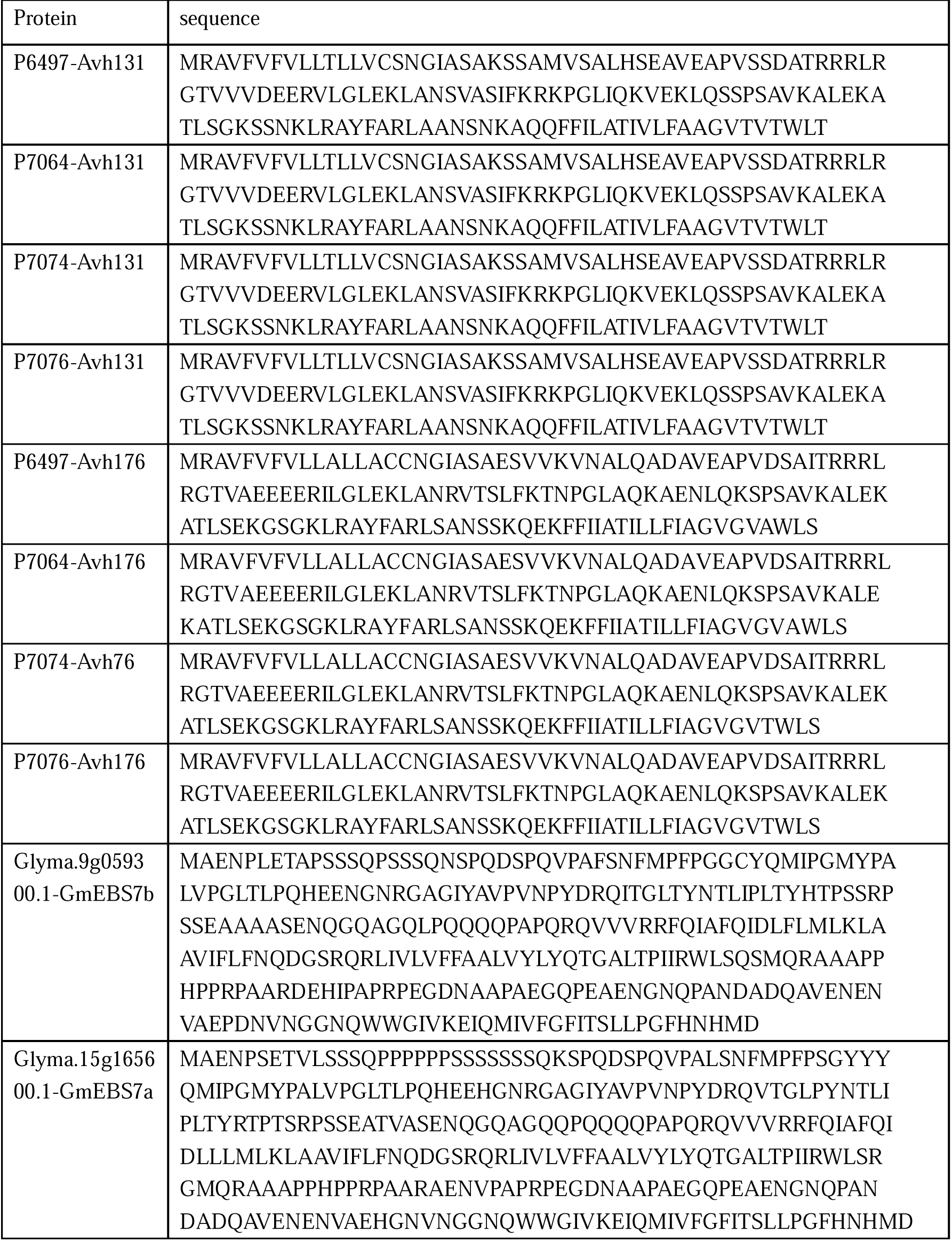
Amino acid sequence in this study.

**Table S2.**
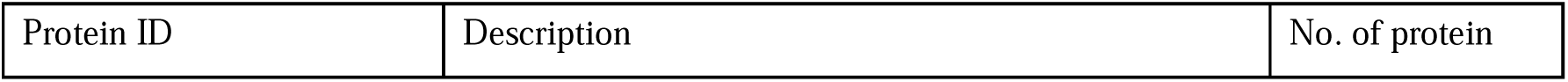
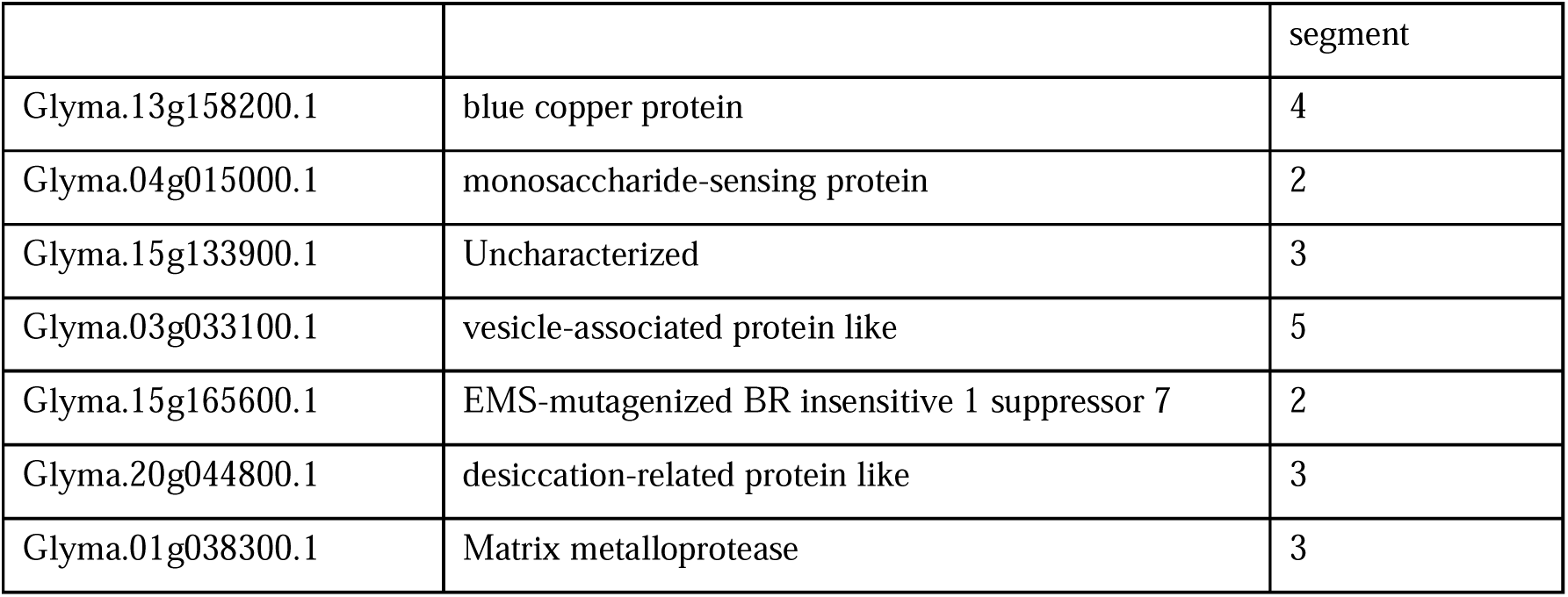
Candidates for host target of Avh131.

**Table S3.**
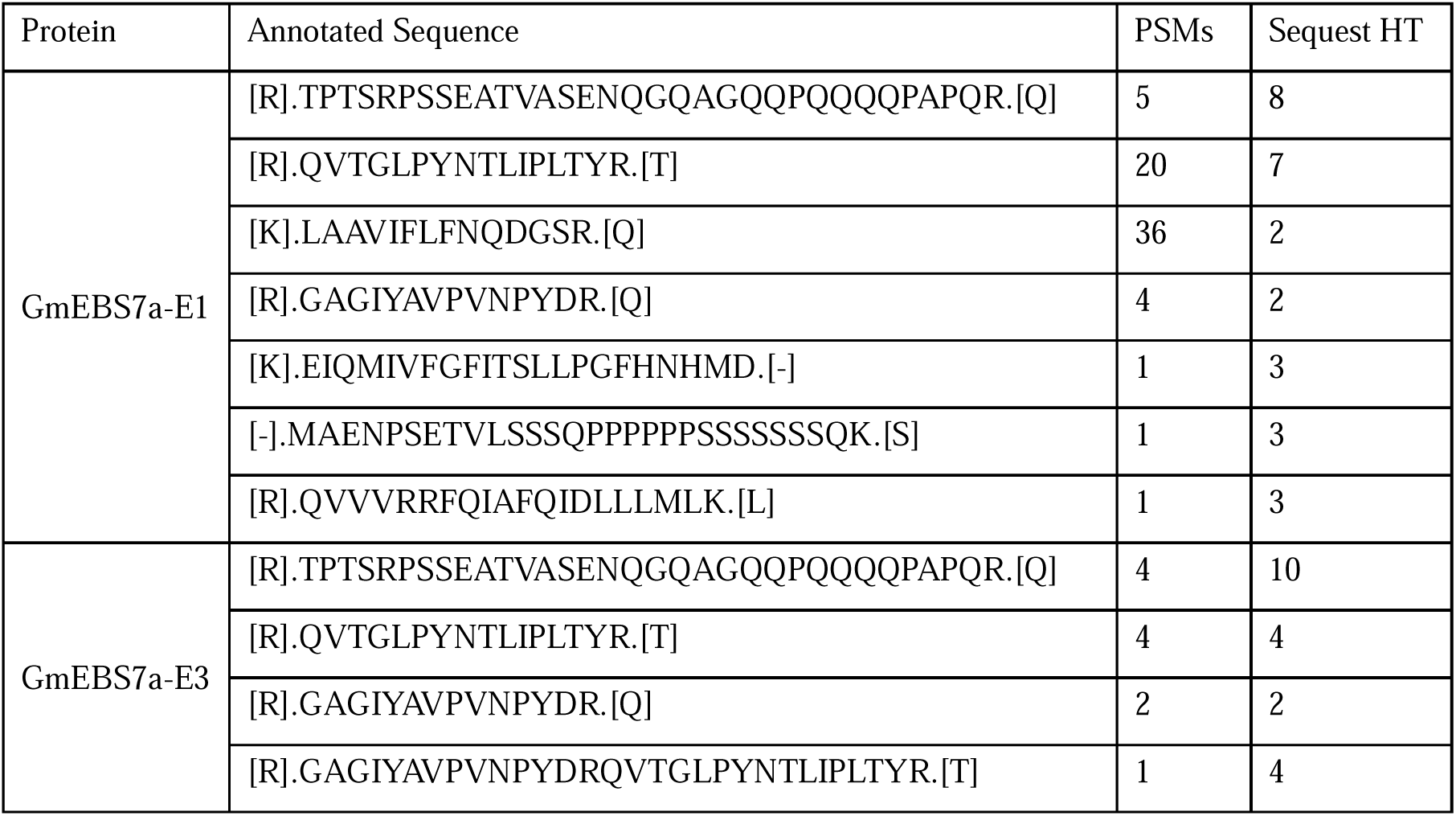
The protein peptide obtained of GFP-GmEBS7a-E1 and GFP-GmEBS7a-E3 by the IP-MS/MS method.

**Table S4.**
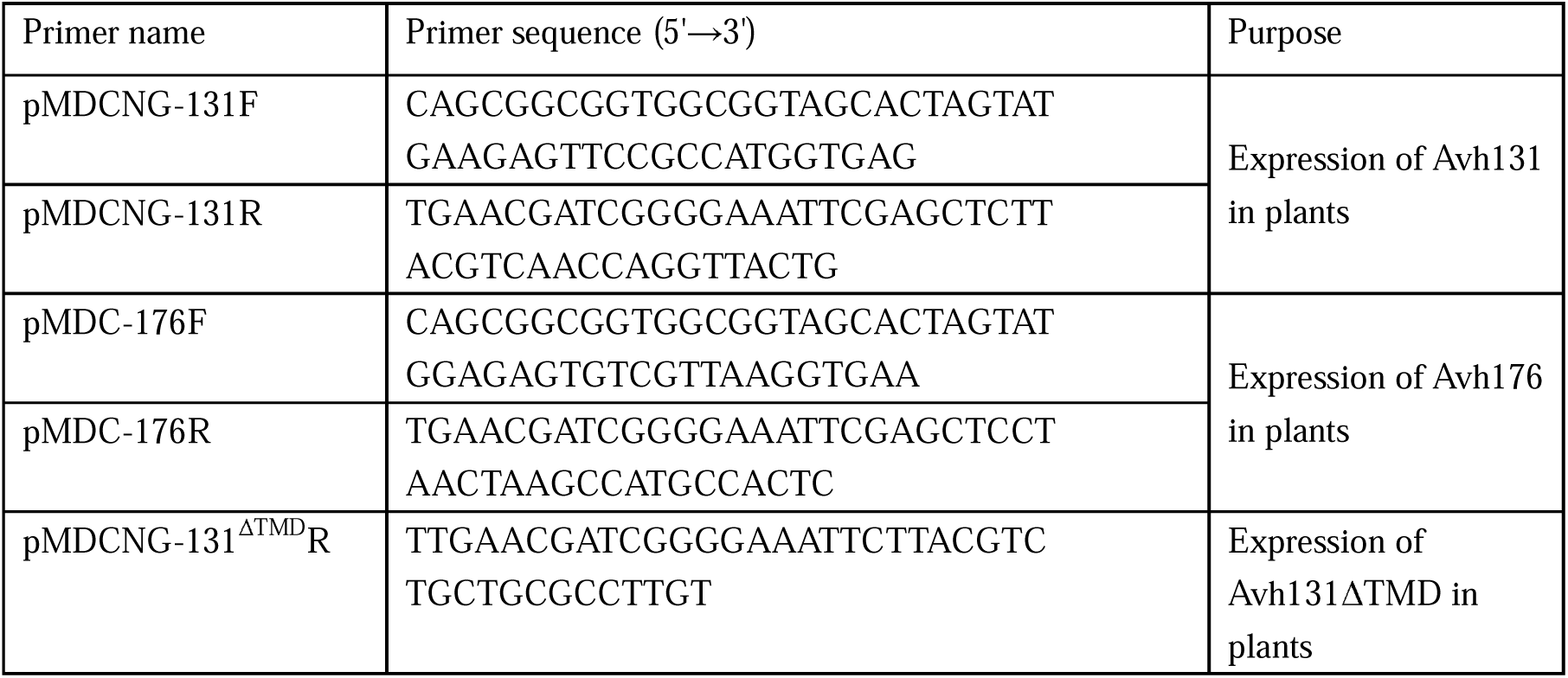
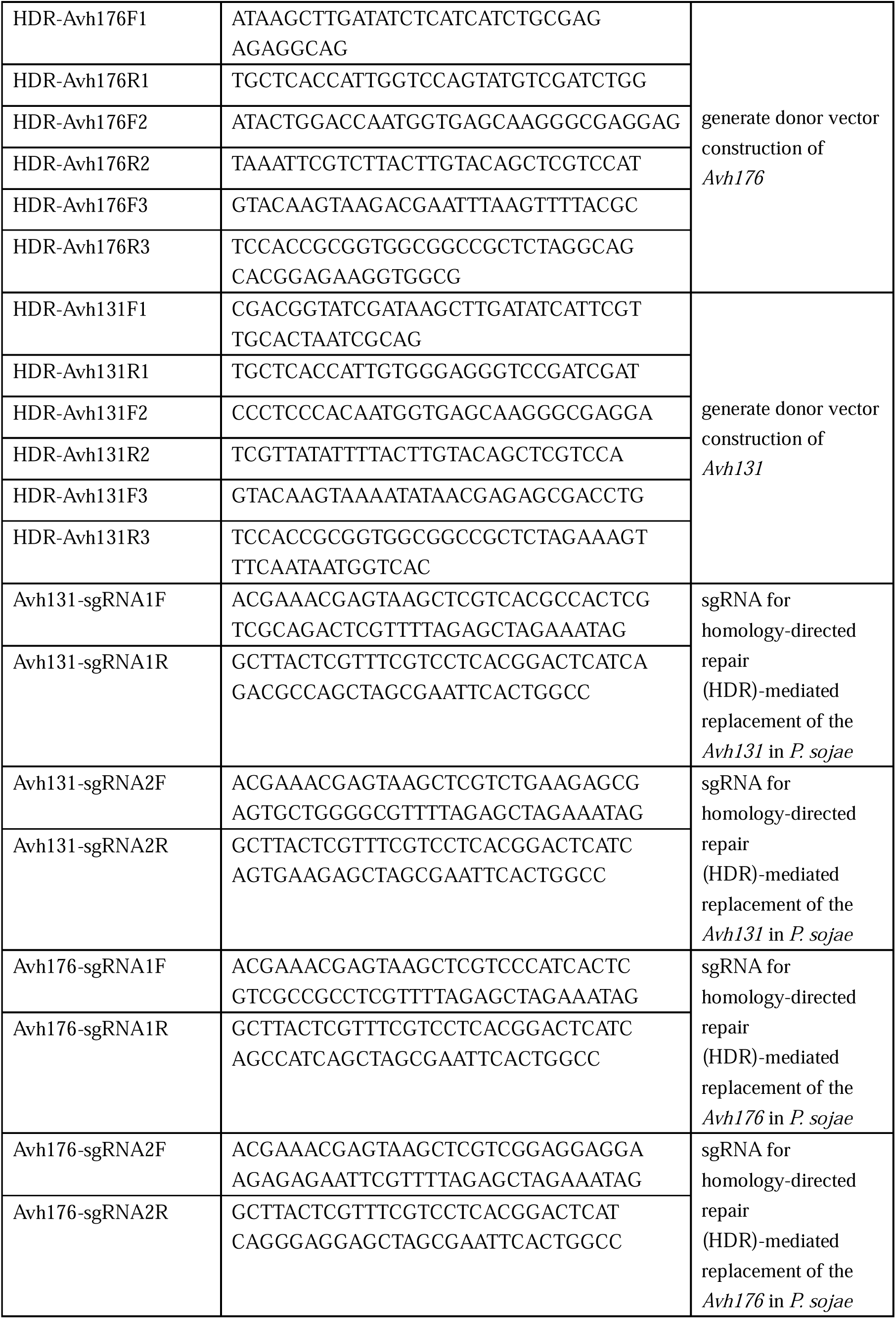
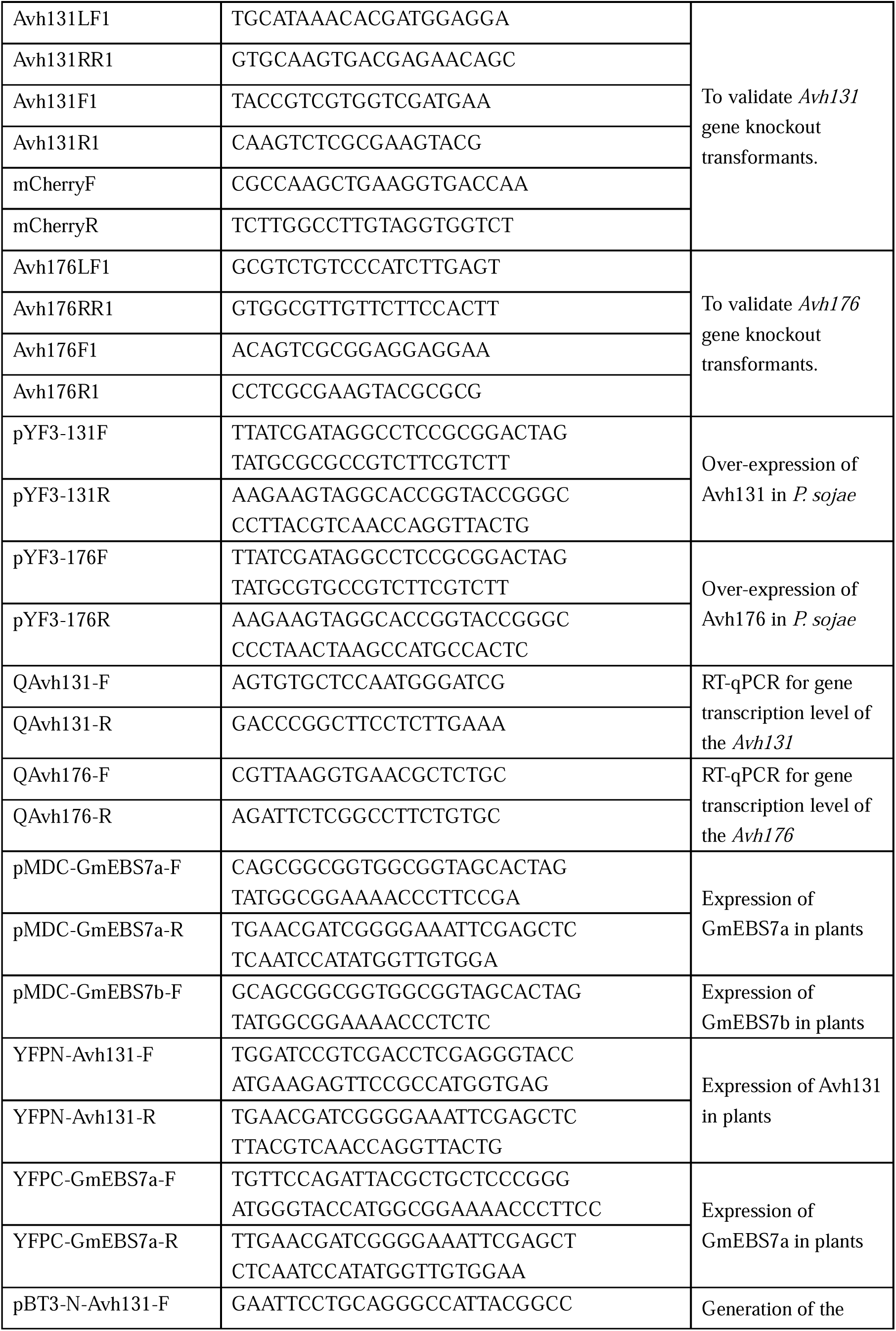
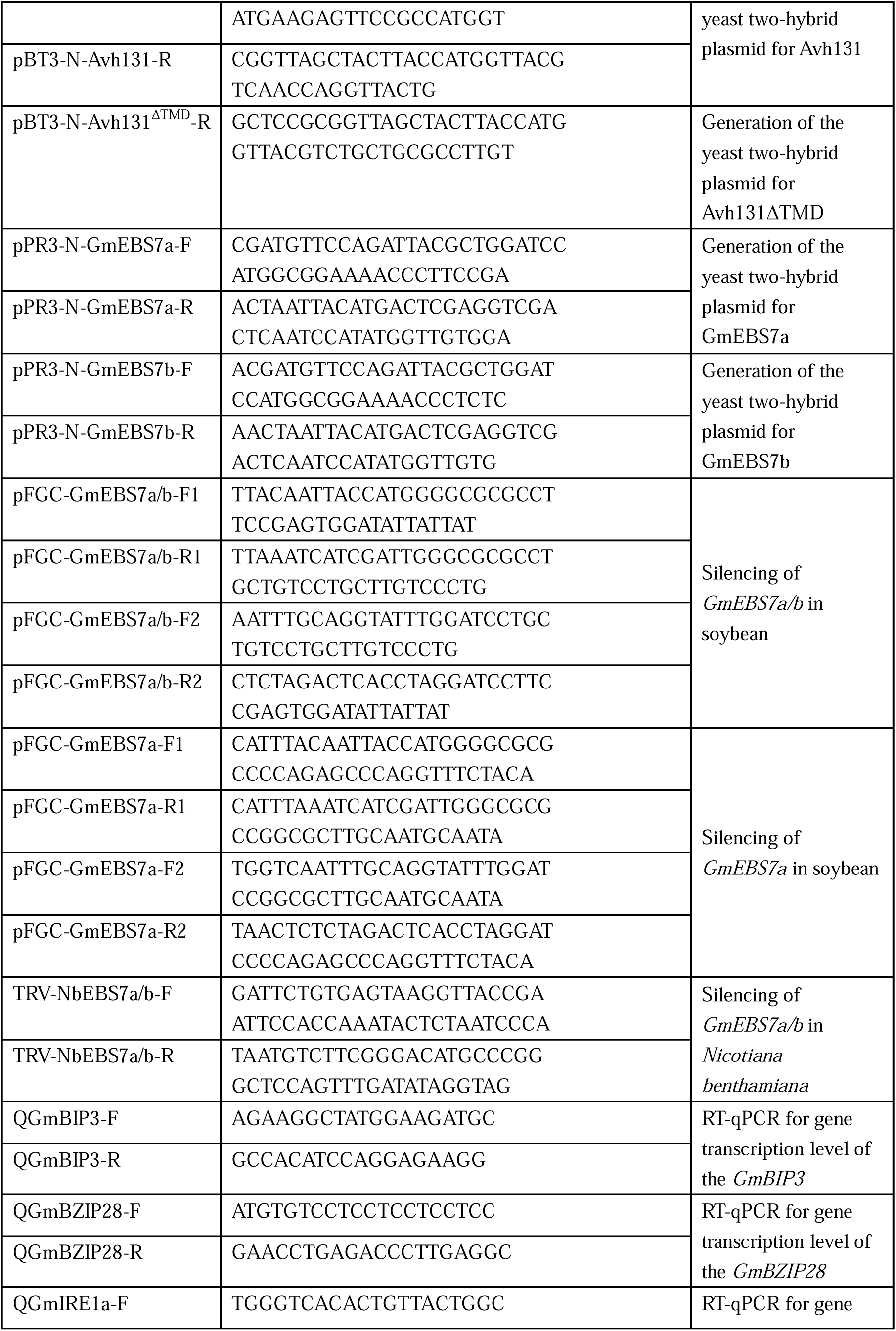
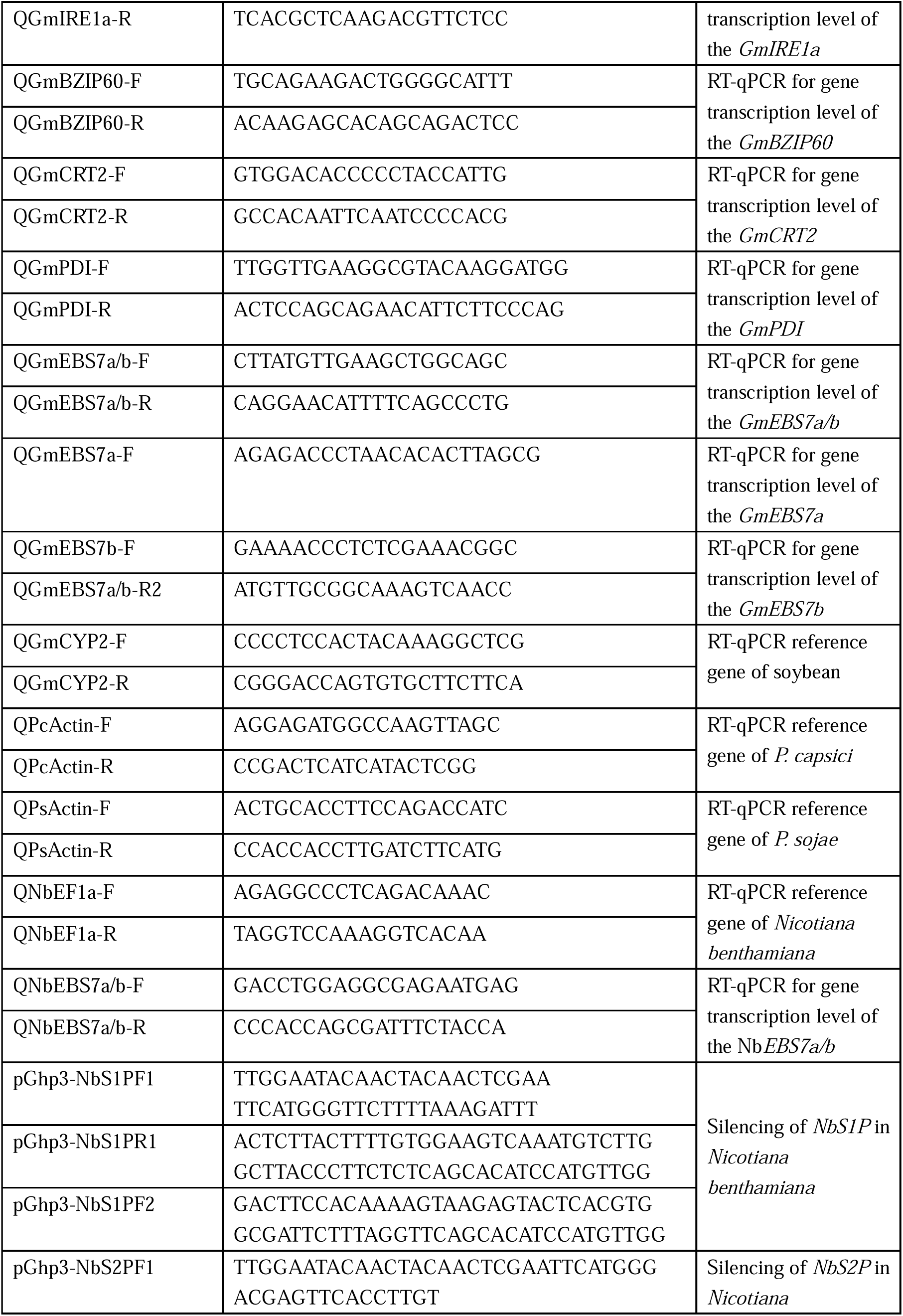
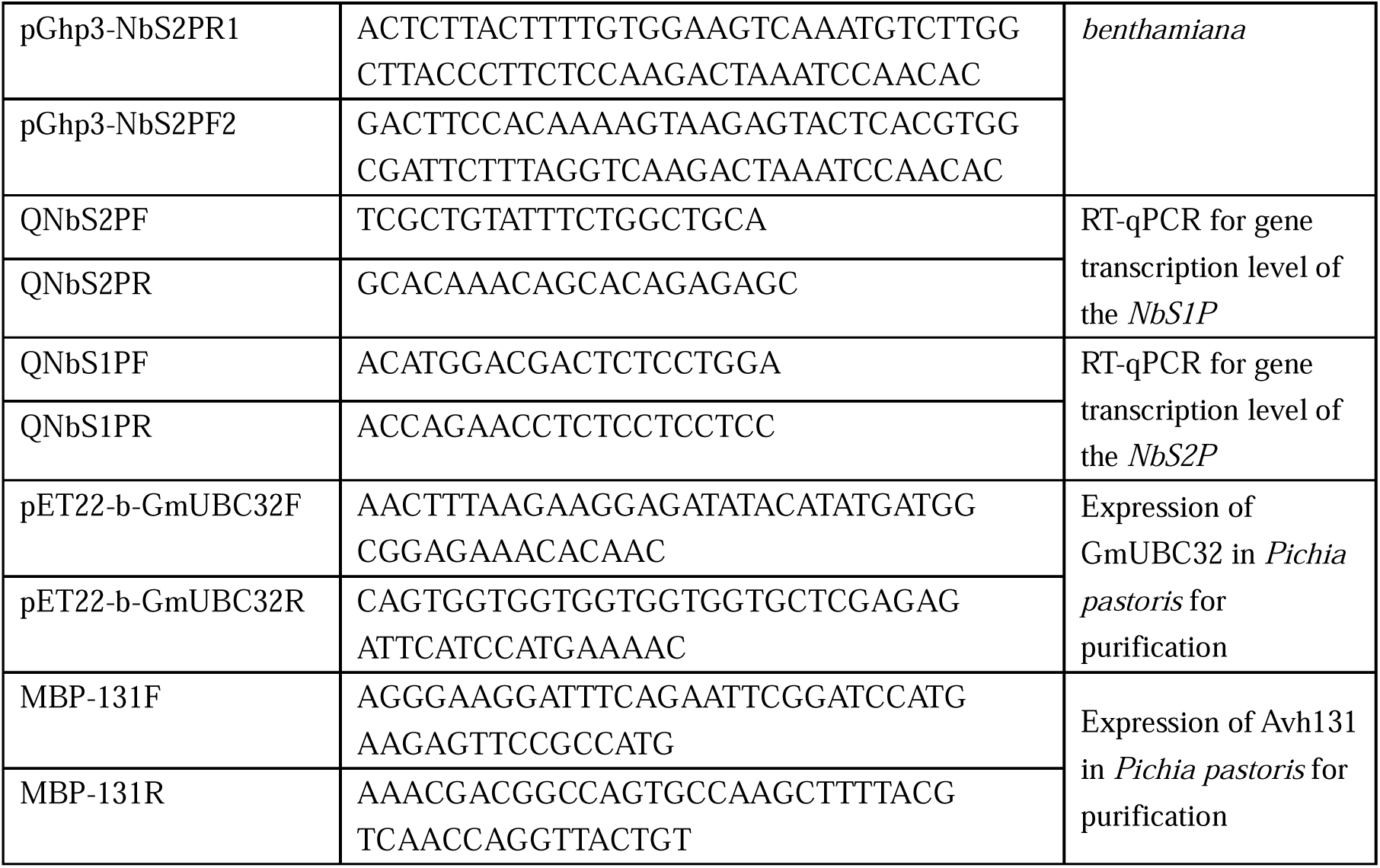
Primers used in this study.

